# Discovering optimal kinetic pathways for self-assembly using automatic differentiation

**DOI:** 10.1101/2023.08.30.555551

**Authors:** Adip Jhaveri, Spencer Loggia, Yian Qian, Margaret E. Johnson

**Affiliations:** TC Jenkins Department of Biophysics, Johns Hopkins University, 3400 N Charles St, Baltimore, MD 21218

## Abstract

During self-assembly of macromolecules ranging from ribosomes to viral capsids, the formation of long-lived intermediates or kinetic traps can dramatically reduce yield of the functional products. Understanding biological mechanisms for avoiding traps and efficiently assembling is essential for designing synthetic assembly systems, but learning optimal solutions requires numerical searches in high-dimensional parameter spaces. Here, we exploit powerful automatic differentiation algorithms commonly employed by deep learning frameworks to optimize physical models of reversible self-assembly, discovering diverse solutions in the space of rate constants for 3-7 subunit complexes. We define two biologically-inspired protocols that prevent kinetic trapping through either internal design of subunit binding kinetics or external design of subunit titration in time. Our third protocol acts to recycle intermediates, mimicking energy-consuming enzymes. Preventative solutions via interface design are the most efficient and scale better with more subunits, but external control via titration or recycling are effective even for poorly evolved binding kinetics. Whilst all protocols can produce good solutions, diverse subunits always helps; these complexes access more efficient solutions when following external control protocols, and are simpler to design for internal control, as molecular interfaces do not need modification during assembly given sufficient variation in dimerization rates. Our results identify universal scaling in the cost of kinetic trapping, and provide multiple strategies for eliminating trapping and maximizing assembly yield across large parameter spaces.

**SIGNIFICANCE:** Macromolecular complexes are frequently composed of diverse subunits. While evolution may favor repeated subunits and symmetry, we show how diversity in subunits generates an expansive parameter space that naturally improves the ‘expressivity’ of self-assembly, much like a deeper neural network. By using automatic differentiation algorithms commonly used in deep learning, we searched these parameter spaces to identify classes of kinetic protocols that mimic biological solutions for productive self-assembly. Our results reveal how high-yield complexes that easily become kinetically trapped in incomplete intermediates can instead be steered by internal design of rate constants or external and active control of subunits to efficiently assemble, exploiting nonequilibrium control of these ubiquitous dynamical systems.

## INTRODUCTION

Self-assembled protein complexes evolved to function not only under selection for specific structures, but under selection for specific kinetics. Disrupted kinetics of virion assembly significantly reduces infectivity(1), and ribosome biogenesis fails without extensive kinetic control(2, 3). Improving the rational design space for assembly kinetics will significantly enhance efforts to optimize assembly function for synthetic biology and drug delivery(4). However, a significant challenge with kinetic selection is the size of the parameter space that can be sampled and the nonlinear dependence of assembly on diverse components. Furthermore, this inherently nonequilibrium process is not only fundamentally influenced by kinetic selection of dominant pathways via A) binding rates (5), but it can be further driven by time-dependent control via B) subunit ‘activation’ (e.g. post-translational modification(6) or co-factor binding(7)) or C) enzymatically-driven dissassembly of intermediates(8, 9). Here, we therefore turn to gradient-based optimization using automatic differentiation (AD), which is similar to backpropagation used in machine learning and is a remarkably efficient and flexible approach for high-dimensional parameter optimization(10). AD does not require analytical gradients, which is a key advantage for our dynamical systems that lack analytical solutions, and recent applications successfully optimized structures(11) and kinetics of self-assembling crystals and small clusters (12). With AD, we find optimal solutions in the largely unexplored parameter space of rate constants that map to the three kinetic control mechanisms [A,B,C] defined above for biological self-assembly of *N*=3-7 subunit complexes.

Our primary goal is to identify and contrast high-yield and efficient assembly pathways, which is challenging due to kinetic trapping. We focus on kinetic traps that occur due to depletion of monomers before intermediates complete growth (13), thus assuming no misassembled intermediates as non-specific interactions (14-16) are eliminated. For single-component assemblies like viruses, the transition to kinetically trapped assembly can be theoretically estimated (17, 18), but the timescales of kinetically trapped systems have not been characterized to our knowledge. By defining timescales here that we demonstrate to exhibit universal scaling with subunit free energies and concentrations, we provide formulas for the cost of kinetic trapping, emphasizing how much is gained via optimization. To avoid traps, one approach is to identify an optimal subunit-subunit interaction strength(19, 20). However, for viruses (7, 20) and reversible heteromeric assemblies(21), this optimal energy occurs over a narrow range that shrinks with increasing assembly size(19, 22), and identifying this optimum cannot be done *a priori* (17, 21). Trapping can be alternatively avoided for distinct subunits by either introducing variations in subunit energetics(22, 23), or by variation in stoichiometries of monomer subunits(14). Subunit diversity also allows for improved control over equilibrium assembly yield by selecting against mis-assembled(14) or morphologically variant complexes(24). Here we consider variations in rate constants as a distinct strategy for avoiding traps via both internal or external and active control of subunits. While recent work quantified how slowing dimerization rates relative to higher-order reactions could dramatically improve efficiency (21), the parameter space was limited to a single rate constant. With our results, we systematically explore large parameter spaces of kinetic control with AD, showing how subunit diversity can improve designability of self-assembly without compromising efficiency.

## Models and Optimization

In our approach we 1) set up a differential equation-based model of a self-assembly topology with initial concentrations and rate constants. 2) Numerically integrate these ordinary differential equations (ODEs) to a predefined time *t*_stop_. To use AD routines in pyTorch, we implemented a differentiable numerical integration algorithm. During integration we store variables and partial derivatives at each step for AD to construct the gradient of our objective function *L* following the chain rule under any assembly protocol. The objective function is the yield of completed complexes at *t*_stop_ plus regularization terms to constrain our rates to physical regimes (Table S1). 3) Modify our rate parameters *k* by following the gradients 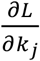 that we automatically constructed, keeping free energies unchanged. 4) Return to step (2) and iterate until we reach the optimal yield by *t*_stop_. We verified our results are not sensitive to our selection of *t*_stop_ (Fig S1) or the learning rate that controls convergence (Fig S2). A detailed description of the algorithm and validation is provided in the SI. Our code is open-source via github.com/mjohn218/KineticAssembly_AD

Our three classes of biologically motivated protocols have multiple rate-constants free for optimization, but vary in their ease of molecular designability or external control (Fig 1). In protocol A, we modify pairwise binding rates, thus reflecting optimal internal design of the molecular interactions that drive assembly. In our ‘rate growth’ model (A1), we allow association rates to accelerate as assemblies grow, which biophysically must reflect cooperativity(25), conformational changes(26), or active modification of interfaces(27). Dimerization reactions occur with the same rate, *k*_2_, trimerization with the same rate *k*_3,_ etc, so this model can apply to both homo- or hetero-subunit assembly. With *N* subunits there are *N*-1 free rates to optimize. In our ‘diversification’ model (A2), we allow independent binding rates between distinct dimers (e.g. *k*_12_ ≠ *k*_23_), reflecting heterogeneity in binding interfaces. We constrain all higher-order steps by the dimer rates of participating interfaces, implying that no molecular modifications are necessary as intermediates grow (Fig 1), giving *N*\*(*N*-1)/2 dimerization rates to optimize. In protocol B, we introduce external control via titration of individual subunits with distinct titration rates for each type of subunit, effectively modifying the initial conditions. Titration can be interpreted as controlling the physical appearance of subunits (e.g. via *in vivo* translation or *in vitro* titration), or as a rate of activating subunits into assembly-competent conformations. We contrast a multi-rate model with *N* distinct rates for all subunits *α*_1_, *α*_2, …_ *α*_N,_ versus a single-rate model with *α* for all subunits, as that model applies to homo-subunit assemblies. Subunit titration is stopped once it reaches the target concentration.

**Fig 1.**
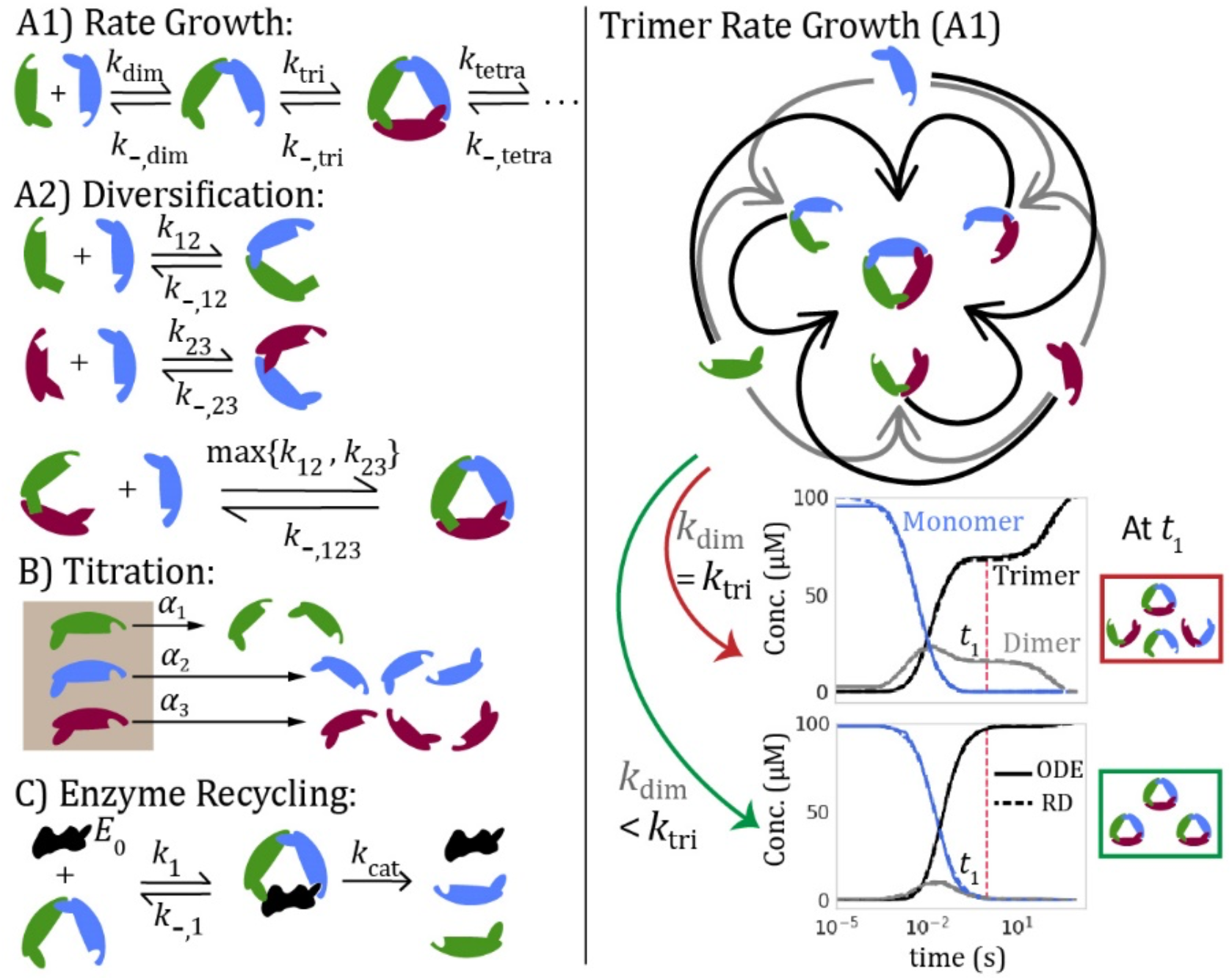
Description of kinetic protocols optimized to achieve efficient assembly. In the left column we illustrate the three kinetic protocols that we contrast to prevent (A and B) or correct (C) kinetic trapping. In A1 and A2 we optimized pairwise binding rates between subunits, in B we titrate in subunits at distinct rates, and in C we actively disassemble intermediates with an enzyme to recycle monomer subunits. On the right, we show a graphical representation of trimer assembly and the time-dependent yield with two different sets of rates. For the over-stabilized system simulated here (Δ*G*<Δ*G*_opt_), equal rates of dimer and trimer binding result in kinetic trapping as monomers are completely used up in both dimers and trimers (top plot). The plateau at time t_1_ emerges due to the wait for dimers to dissociate. In the optimized rate- growth model (A1), we slowed down the dimer rate relative to the trimer rate to efficiently achieve high-yield by t_1_ (bottom plot). Each of the two scenarios of trimer assembly are also simulated using a spatial stochastic reaction-diffusion method (RD: dashed lines), with excellent agreement.

In protocol C we recycle trapped intermediates via their irreversible dissociation by an enzyme, returning monomers into the pool (*S*_1_*S*_2_ + *E* ⇌ *ES*_1_*S*_2_ → *E* + *S*_1_ + *S*_2_). We evaluate enzyme intervention via a) Single substrate, where the enzyme reacts with only one intermediate and b) Multi substrate, where the enzyme reacts with multiple intermediates but only one of each size (2≤ intermediate< *N*). We optimize the enzyme concentration *[E]*_0_ and association rate of substrate binding (*k*_1_) and catalysis rate (*k*_cat_) for each substrate while keeping the ΔG fixed, giving maximally 2*(*N*-2) + 1 parameters. Protocols B and C both provide time-dependent quality control rather than pre-optimized design. These solutions are critical in cell biology, as *in vitro* reconstitution has shown that simply mixing subunits at time zero often fails for multi-component self-assembly (28-30)

We contrast two limiting assembly topologies, the fully connected graph where all subunits contact one another, and the ring topology where a single cycle incorporates all subunits. We exclude linear topologies as they have zero cycles and do not become trapped with equal stoichiometries, and we stop at size *N*=7 to focus on more compact structures where the fully connected topology offers a reasonable limit on subunit interactions. All intermediates can combine if sterically possible (i.e. dimer+dimer*⇌*tetramer), but for rate-growth we prohibit these non-monomer growth pathways. To focus on rate variation, we use equal concentrations of subunits and equal free energies for all interactions. We assume no cooperativity, and thus the free energies sum across bonds with a stability for *m* bonds given by:

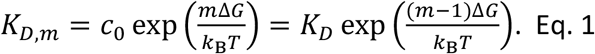

where 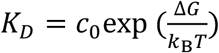, *c*_0_ is the standard state concentration 1M and Δ*G* = *G*_bound_ – *G*_unb_.

We use *K_D_* and Δ*G* interchangeably given they are a log transform apart. We bound *k*_f_ due to the limits of diffusion and the measured kinetics of protein-protein association at 10 or 1*μ*M^-1^s^-1^ (31). The formation of additional bonds during association must therefore slow the off-rates, which become:

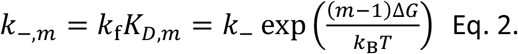

where *k* _−_ is the off-rate for a single bond and 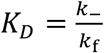. We reproduce the assembly kinetics using structure-resolved reaction-diffusion simulations(32), where association involving two bonds (i.e. dimer+monomer) proceeds in two steps unlike the 1-step model used for the deterministic model (SI and Table S2). By dividing those association rates by two, we recover essentially identical kinetics (Fig 1).

## RESULTS

### With uniform rates, kinetic trapping emerges for systems with maximal yield

In our models, kinetic trapping first emerges for *N*=3 (*N*=2 lacks intermediates). The system becomes starved of monomers before intermediates (dimers) can complete assembly (Fig 1, Fig 2a). To derive expressions that quantify the effective time-cost of kinetic trapping as we vary assembly parameters, we define a trapping factor: TF = τ_2_/τ_1_. The two timescales τ_1_ and τ_2_ delineate the entry into and exit from the trapped regime where the time dependence of the yield (e.g. trimers(*t*)/trimers_MAX_) is stationary (Fig 2a). During the initial growth phase free monomers are abundant and the target complex forms along with intermediates, whereas during the delayed later growth phase(s), monomers are consumed and the target complex grows only as existing intermediates dissociate. Thus TF ≥ 1, with equality when no trapping occurs.

**Fig 2.**
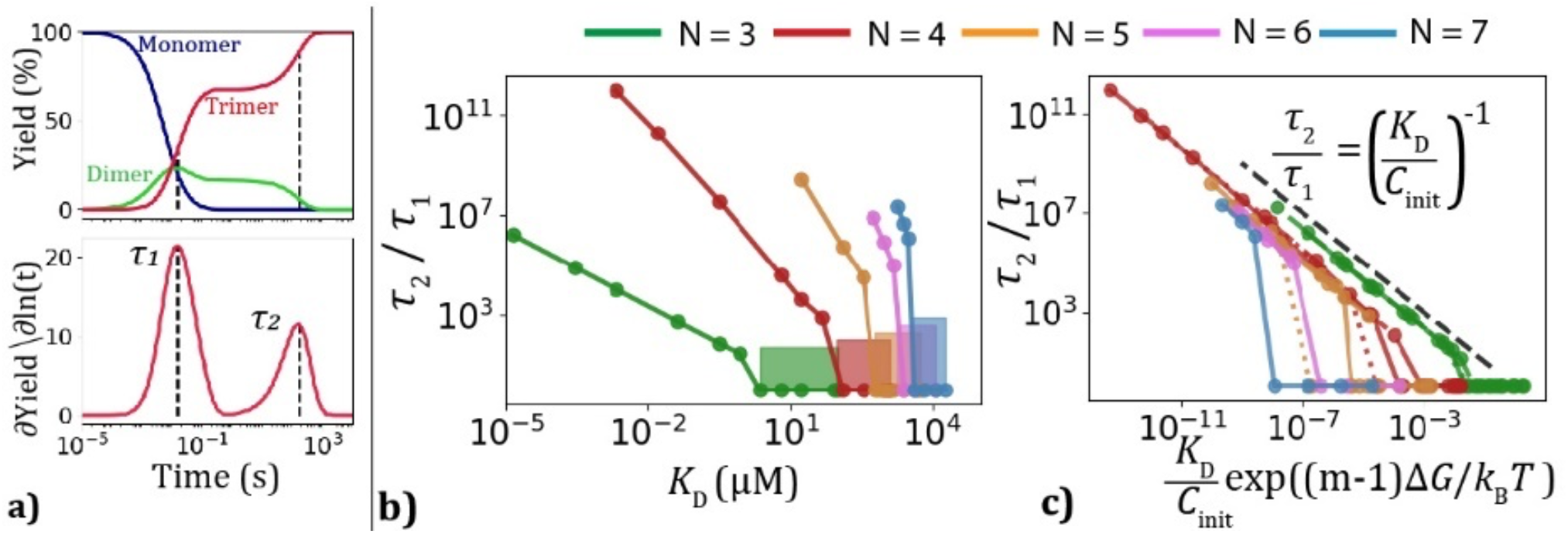
The time-cost of kinetic trapping quantified by the trapping factor. a) Time-dependence of trimer formation for Δ*G* = -20 *k*_B_*T* (*K*_D_=2x10^-3^ μM) and *C*_init_=100 μM reveals a trapping plateau vs ln(t). The derivative of assembly yield with respect to ln(*t*) identifies two maxima that separate entry τ_1_ and exit τ_2_ from the stationary region, which we use to define the trapping factor TF=τ_2_/τ_1_. b) TF increases with stabilizing *ΔG*(K_D_) and scales worse with increasing N=3 (green) to N=7 (blue). The shaded boxes represent the *ΔG* window for each assembly size where yield is >95% and TF=1. This narrow window shrinks as expected with increasing *N*, and encompasses *ΔG*_opt_ where efficiency for equal rates is fastest. *C*_init_=100 μM c) The TF shows power-law scaling with exponent *γ*=-1 (dashed black) vs the stability of the largest intermediate as *N* or *C*_init_ are varied, where *m*=*N*-2. Dashed vs solid colors are distinct values of *C*_init_. We can derive values for the x-intercept vs *N* or *C*_init_ by assuming TF→1 as *ΔG*→*ΔG*_opt_. The power-law scaling of the TF does not fully extend to the line where TF(Δ*G*)=1, showing a drop-off. This is because for a small range of Δ*G* < Δ*G*_opt_, we still measure TF=1 when two distinct peaks have not yet emerged in the ln(*t*) kinetics (a) despite slowed assembly.

Trapping only occurs in highly stable systems, as an equilibrium monomer pool emerges when yield drops below ∼99% with weakening Δ*G* (Fig S3, S4). With equal rates for all binding reactions, efficiency as evaluated by time taken to reach 95% yield, τ_95_, is fastest at Δ*G*_opt_ just before trapping sets in, and the TF sharply increases from 1 around this transition in yield Δ*G*<Δ*G*_opt_ (Fig 2 and Fig S4). Trapping is problematic not only due to efficiency but also the trapped yield (22), which drops from 67.39% for the trimer (see SI for derivation) to ∼55% for the tetramer and only ∼20% for the 7-mer (Fig S5). Accelerating all rates uniformly shortens τ_95_ but does not alter the TF; faster rates speed up the timescales to both enter (τ_2_) and exit (τ_2_) the trapped regime, rather than eliminating it (Fig S6), which can be proved by renormalizing time by *k*_f_*C*_init_ (SI).

### The trapping factor shows a nearly universal dependence on *N* and ΔG

Timescales once trapping has set in have not been previously quantified to our knowledge, and here we find that the TF has a universal power-law dependence on *N*, Δ*G*, and *C*_init_ (Fig 2). The entry time τ_1_ is driven by the speed of association *k*_f_*C*_init_ causing an increasing TF with concentration as the exit time τ_2_ is unaffected (Fig S4). The exit time τ_2_ is dominated by the lifetime of the most stable intermediate, which has a dissociation rate 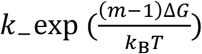, with *m* the number of bonds broken for a monomer to dissociate from this intermediate (e.g. *m* = *N* − 2). The ratio of these timescales is then (Fig 2c):

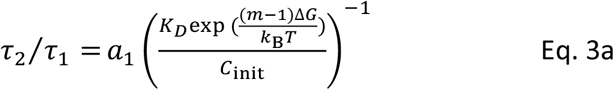

and the positive dimensionless constant *a*_*%*_ is relatively well-predicted by the free energy when the efficiency is fastest Δ*G*_opt_ (see Table S3), which varies with number of subunits *N* and *C*_init_ giving

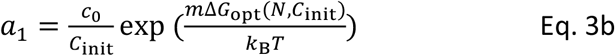

or together, 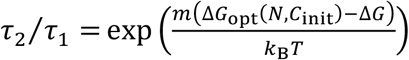. The τ_95_ follows the exit time τ_2_ once the system becomes strongly trapped, becoming independent of entry time τ_1_ and increasing *k*_f_*C*_init_, resulting in the asymptotic scaling relationship (Fig S4)

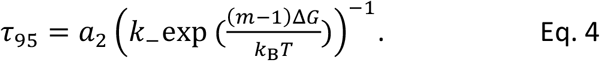

where *a*_2_ is a dimensionless constant that depends on *N* subunits and not *C*_init_. In this strongly trapped regime, the efficiency of assembly cannot be improved by altering initial concentrations, unlike for the weakly or non-trapped systems (Fig S4).

These equations illustrate how larger assemblies with more bonds per subunit will have exponentially worse trapping at identical ΔG values (Fig 2b), as higher order intermediates have much slower dissociation times (Eq. 2). For the ring topology, however, all subunits have only two binding partners and therefore all intermediates have the identical lifetimes, with *m*=1. All ring sizes therefore have a TF scaling with 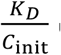 that is now independent of *N* (Fig S7). The trade-off, however, is that without additional stabilizing cycles, larger ring structures need increasingly more stable Δ*G* values to reach high yield, and the efficiency at Δ*G*_opt_ is significantly worse due to the ease of dissociation for all intermediates (Fig S8).

### By slowing down all dimerization reactions relative to higher intermediates, kinetic traps can be eliminated: Rate Growth

Kinetic trapping emerges for highly stable systems, but these results are dependent on our model parameters: uniform rates, initial bulk concentrations (equal), and no external activity on intermediates. Using AD (Methods), we find an optimal pathway to efficiently reach high yield is to create a separation between the dimerization and elongation steps by slowing dimerization, e.g. 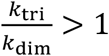 (Fig 3), where we measure normalized efficiency via 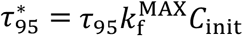, and 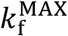 is the fastest of all association rates. A slower dimerization step ensures that monomers are conserved long enough to be incorporated into a fully formed complex (Fig S9). The optimal ratio 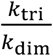 increases with stabilizing ΔG until it reaches a plateau (Fig 3d,f), which we can derive for the trimer system under irreversible assembly (SI). 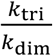 must also increase with *N*, thus driving a reduction in efficiency for larger assemblies (Fig 3d,b). We find the optimal efficiency when rates for growth beyond the dimer all proceed at the same speed (*k* _tri_ = *k* _tetr_ = *k* _pent_ …), but this solution requires the largest value 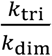 The ratio 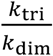 can be significantly reduced if we introduce an additional hierarchy where 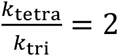, for example, providing flexibility in growth rates with a <2-fold loss in efficiency (Fig S10). Having a smaller separation in rates would require fewer conformational changes to the existing interfaces(25, 26) to speed-up subsequent growth steps. In the ring topology, the optimal ratio between rates is reduced compared to the fully connected topology (Fig 3f), but results in minimal improvement in efficiency (Fig 3b). The optimal set of rates we find for the fully connected topology still works for the ring topology, and for all topologies between the ring and fully-connected we can thus infer that this solution similarly will succeed in evading traps.

**Fig 3.**
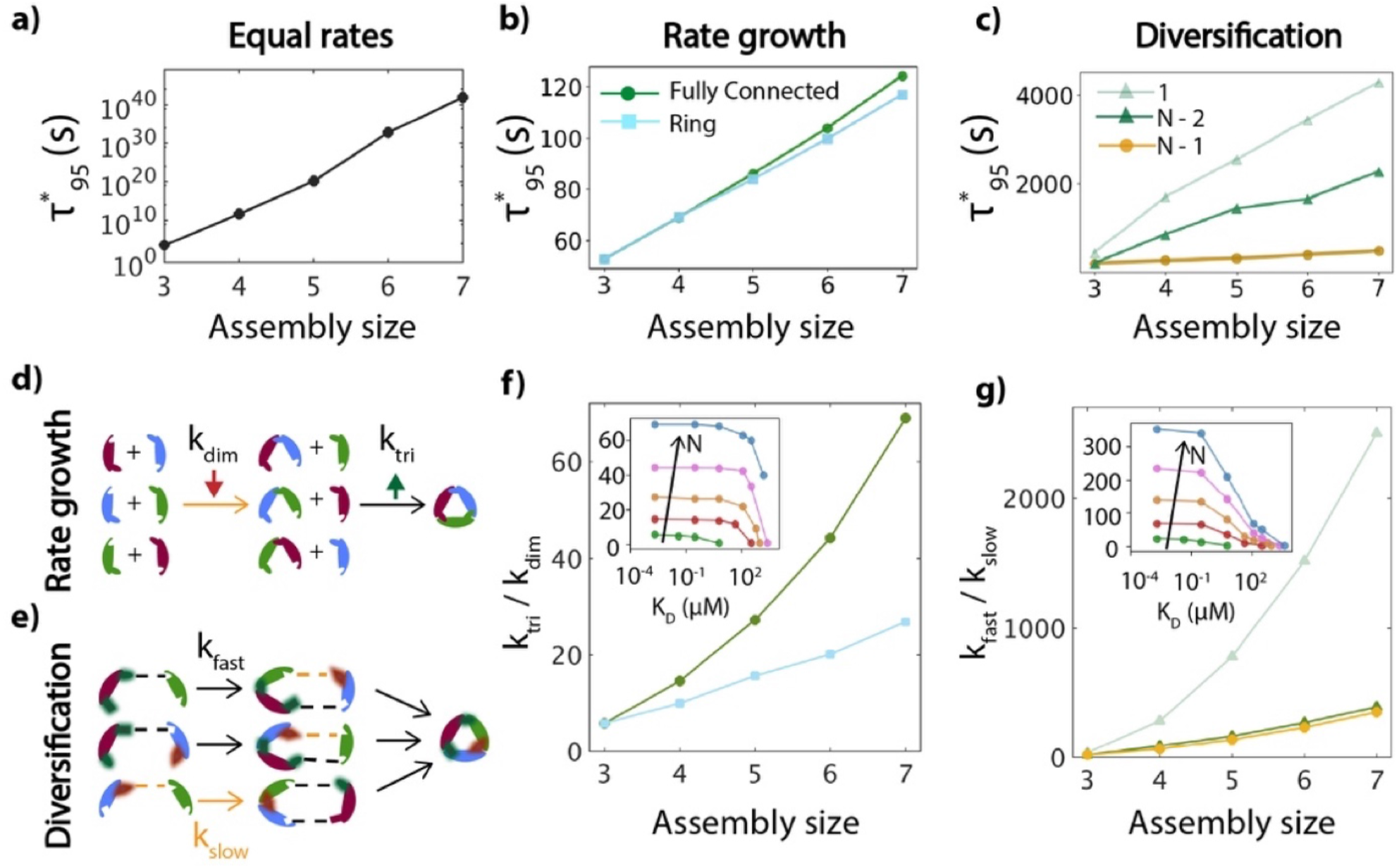
With internal design of binding rates at fixed free energies, efficiency is dramatically improved. a) With equal rates prior to optimization, the timescale 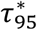 grows exponentially with *N* as predicted by Eq 4, 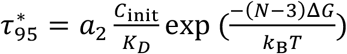. Although *a*_2_ decreases by ∼10^4^ from *N=*3 to 7, this is small compared to the exponential factor which increases by 10^34^. For ease of comparison, an approximate fit of 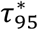 to a power law *N^γ^* gives *γ* ≈90. b) In the rate-growth model, timescales are 1000-fold improved for the trimer and dramatically improved with increasing *N*. A power-law fit gives *γ* ≈0.7. Ring topology (blue) has similar efficiency to fully connected (green). c) For the Diversification model, optimized models with *N*-1 interfaces accelerated (gold) have similar efficiency to the rate-growth, with γ ≈ 1. With *N*-2 (green) or 1 interface (mint) optimized, the efficiency is worse but still a significant improvement over (a). d) Dimerization is slowed relative to the rate of trimerization and subsequent steps. e) If all interfaces on one subunit (red subunit) dimerize faster (*k*_fast_: black dashed) than all other interfaces at *k*_slow_ (gold dashed), traps are avoided. f) The optimal ratios of 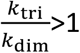 increase with N, and are higher for the fully connected topology. The inset shows how these ratios increase with more stable *K*_D_ until they plateau. Holds for all *N*=3 to 7. g) With dimer diversification, optimal ratios of *k*_fast_ /*k*_slow_ also must increase if fewer interfaces can be accelerated (mint). Insets shows same plateau behavior in *k*_fast_ /*k*_slow_ as *K*_D_ stabilizes 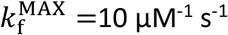, *C*_init_=100*μ*M, *K*_D_=2x10^-3^μM (Δ*G* = −20*k* _B_*T*).

### By slowing down some of the dimerization reactions, kinetic traps can be eliminated

Using AD, we find a second way to evade trapping is to create diversification between dimerization reactions, while all higher-order assembly steps are constrained by these same dimerization rates (Fig 3c). As some dimers form slowly relative to others, this ensures that a subset of monomers remain in the system long enough to complete assembly of fast-forming intermediates. For optimal efficiency, one subunit engages in fast binding interactions while other subunits have slower binding (Fig 3e). Similar to the rate-growth model, with increasing stability or *N* subunits, the separation between the fast and slow rates must increase to avoid traps, but with ∼10 fold more variation than in the rate-growth model (Fig 3g).

Because our optimal solution requires that all the *N*-1 interfaces on one of the subunits must have fast binding, we considered more ‘designable’ solutions where a smaller subset of interfaces (*n*<N-1) must be accelerated (Fig S11). These solutions are not as efficient given that fewer binding interactions are fast (Fig 3c), but they are nonetheless a vast improvement over uniform rate efficiency. While we can design fewer interfaces (even 1) to have an accelerated rate, the ratio of these rates must be more extreme to avoid trapping (Fig 3g). Diverse subunits again helps, with improved efficiency when a hierarchy of rates (not just *k*_fast_ and *k*_slow_) are distributed across interfaces (Fig S11).

### By externally controlling titration rates of subunits, kinetic traps can be eliminated

Instead of modifying internal subunit binding rates, here we keep those rates fixed at nonoptimal equal values that lead to kinetic traps, but introduce titration rates (α_1_, α_2_, α_3_, …) to control subunit concentration in time (Fig 4). Titrating too quickly leads to more bulk-like behavior and fails to eliminate traps, whereas slower titration delays assembly (Fig S12). The optimal rate decreases as ΔG stabilizes (before plateauing-Fig 4) because dissociation of dimers helps replenish the monomer supply over the timescales of titration. Optimal rates are faster with higher subunit concentrations or faster binding rates because individual assemblies form more quickly (Fig 4b). Following recent work (7) we can show that for a single titration rate (SI), the optimal value should scale as

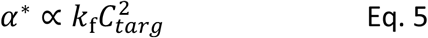

and we found this same scaling numerically through our optimization approach (Fig 4c).

**Fig 4.**
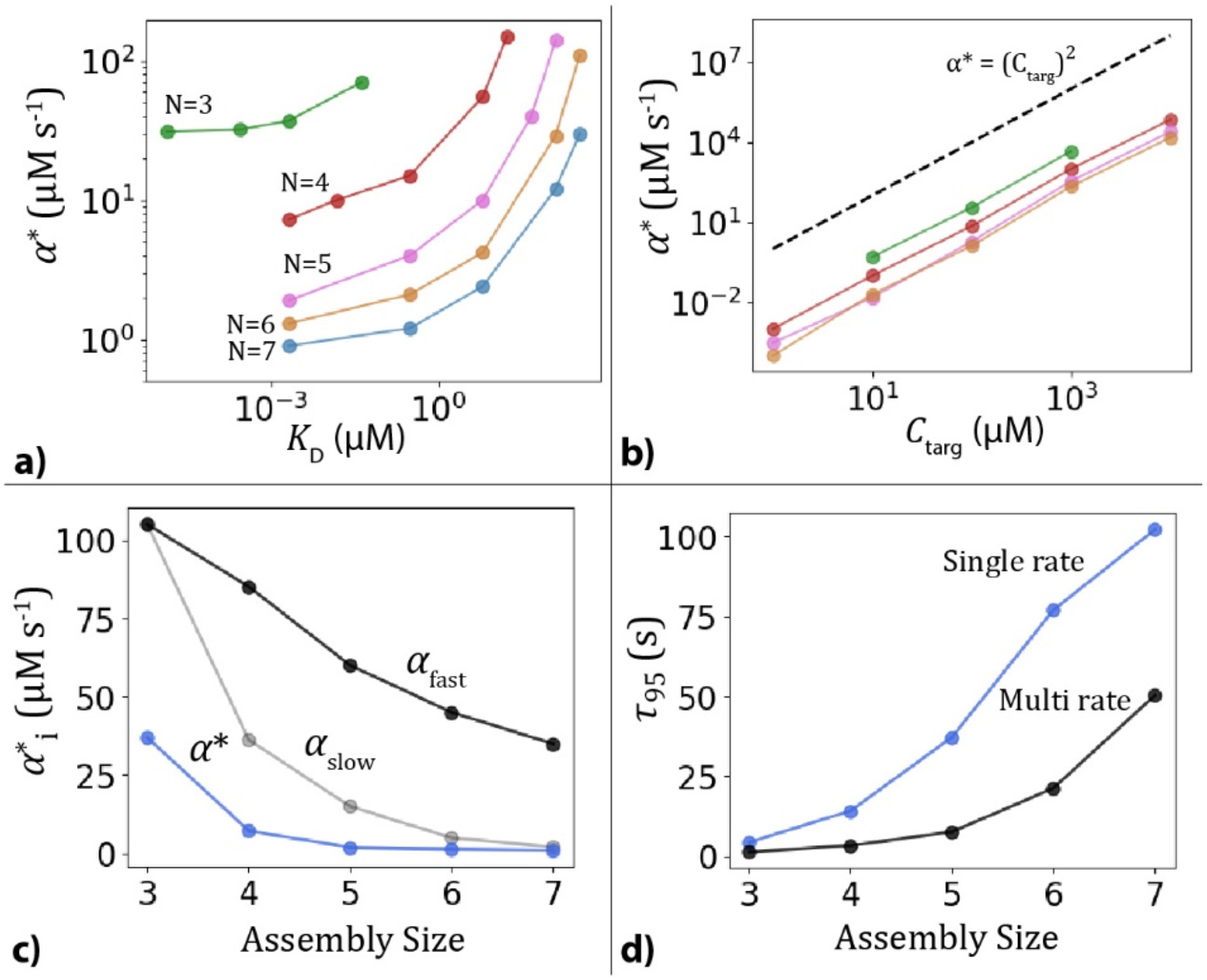
Optimized titration rates are more sensitive to thermodynamic parameters of the trapped system and benefit more from diverse subunits. a) The optimal titration rate α* (single-rate) slows until it plateaus as the ΔG stabilizes (lower *K*_D_). Fastest α* is for when trapping first sets in (ΔG<ΔG_opt_). b) The optimal titration rate (α*) can speed up with increased *C*_targ_, as *C*_targ_^2^ (Eq. 5, dashed black line). Colors correspond to *N* subunits as labeled in (a). c) When each subunit is titrated at a distinct rate, the optimal titration rates follow a hierarchy (from *α*_fast_ (black) to *α*_slow_(gray)) which allows for faster titration as compared to the single rate (*α*\*-blue). d) With faster titration possible for the multi-rate model (black line), efficiency is improved over single-rate (blue); the multi-rate can only be implemented if subunits are distinct. Fully connected topologies.

Although the single-rate scheme works for all *N* (including >3000 subunits(7)), the multi- rate scheme always improves efficiency (Fig 4f). With diverse subunits in the assembly, two subunits can be present in the bulk (α_1_*=∞, α_2_*=∞), with the remaining components activated over a hierarchy of rates that are all faster than α^∗^ from the single-rate scheme (Fig 4). The efficiency gain from using the multi-rate hierarchy increases with *N*, underlining how additional parameters improves control over the assembly process. Lastly, we find that if we are willing to accept partial trapping and yield reduced from 95 to 90%, we can speed-up titration and noticeably improve efficiency (Fig S13).

### Trapping can be corrected by recycling monomers using enzymes

In this protocol, we do not try to prevent trapping, but instead perform quality control by an ATP-consuming enzyme. By recycling subunits from specific incomplete intermediates (the substrate), the enzyme amplifies completion of other intermediates to improve yield per time (Fig 5). When optimizing the enzyme to dissociate only a single substrate, the substrate must be an *N*-1 sized intermediate to eliminate traps. The enzyme must bind to its substrate faster than the competing subunit for effective recycling of monomers, accelerating *k*_1_[E_0_] (Fig S14). The overall assembly efficiency is then limited by *k*_cat_, which cannot be too fast because it must balance recycling of subunits against availability of other intermediates to prevent immediate reassembly of the substrate (Fig 5). For a 7-mer or larger, this single-substrate mechanism fails as the relative concentration of subunits recycled by a single *N*-1 intermediate decreases. By allowing the enzyme to react with multiple substrates (Fig 5), the optimal *k*_cat_ values for distinct substrates can operate faster than the single-substrate value, requiring less delay to release monomers and accelerating assembly formation (Fig 5b).

**Fig 5.**
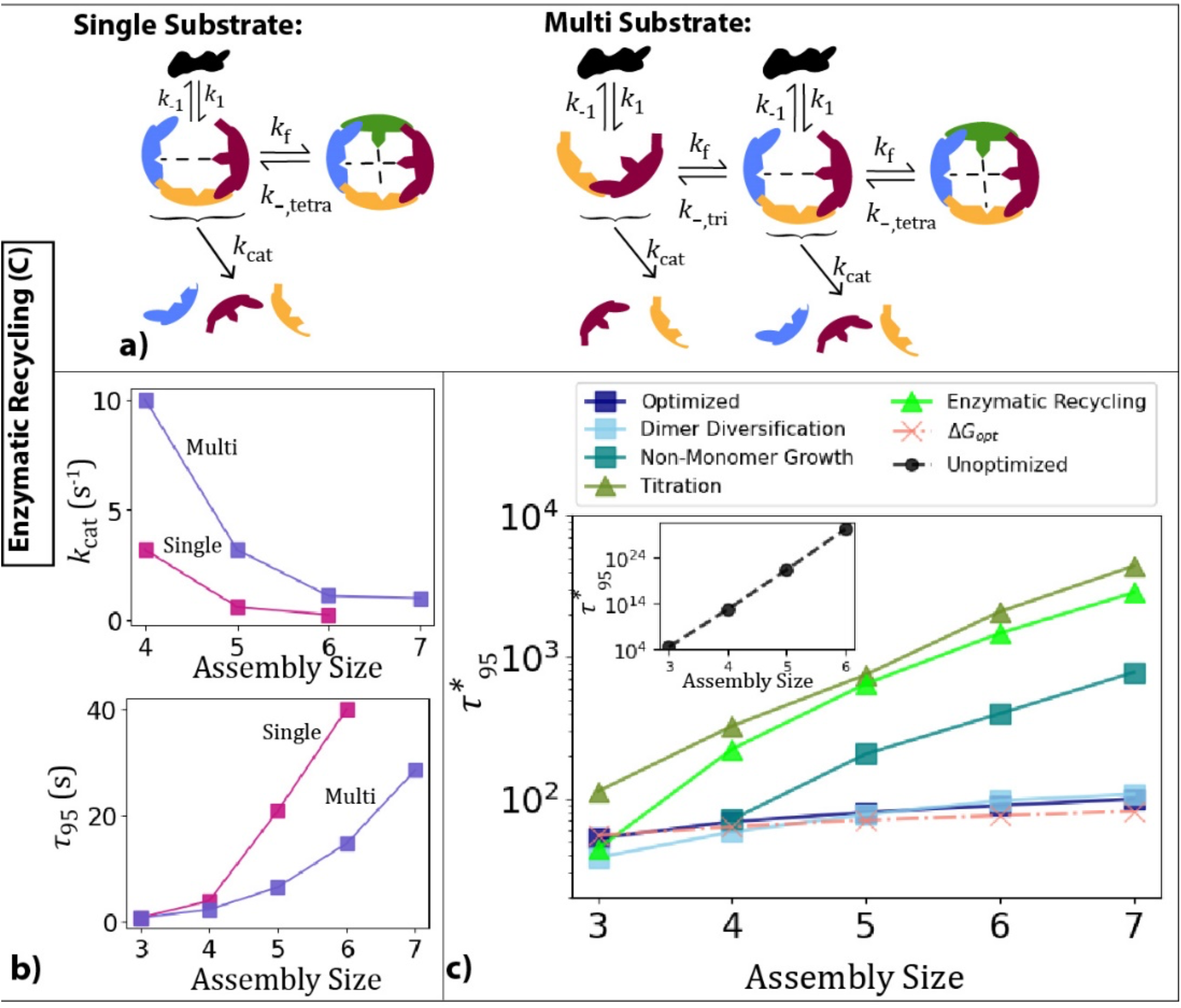
Enzymatic recycling will improve efficiency but not as well as internal rate control. a) The enzyme competes with a particular subunit (green) and binds to a single *N*-1 intermediate that lacks this subunit (Single substrate) or all intermediates that lack this subunit (Multi substrate). All results for fully-connected topology. b) For efficient assembly, there is an optimal rate to dissociate the encounter complex (*k*_cat_) that decreases with assembly size for both modes of recycling. The trimer is an exception, with efficiency increasing monotonically with *k*_cat_ . We show the value that first reaches 95% yield. With multiple substrates, intermediates can be disassembled faster leading to shorter assembly times. For all systems, *ΔG* = -20 *k*_B_*T*, *C*_init_ = 100 μM, *k*_f_ = 1 μM^-1^s^-1^. c) Optimal (normalized) timescales compared for all protocols shows orders of magnitude speed-up with respect to assemblies with equal rates (black dashed, *k*_f_ = 1 μM^-1^ s^- 1^). *ΔG* = -20 *k*_B_*T, C*_init_ = 100 μM. We show efficiency at *ΔG*_opt_ (pink dashed) prior to trapping onset where *τ*_95_ is fastest given uniform rates. For *N=*3-7 we numerically identify *ΔG*_opt_ (Table S3).

## DISCUSSION

In Fig 5 we contrast how highly stable systems that become kinetically trapped with uniform rates achieve order-of-magnitude improvement in efficiency via our kinetic selection of assembly pathways, thus dramatically expanding regimes where high-yield and efficient assembly are met for all *N*. Counterintuitively, slowing down some association rate constants is therefore essential for efficiency. Although our kinetic protocols do not typically exceed the efficiency under the narrow but optimal thermodynamic conditions for the fully connected topology (Fig 5), they do not rely on reversibility and unbinding to prevent trapping (13, 33), but work even in the limit of irreversible binding (Fig S15).

Diverse subunits provide a critical advantage by expanding the parameter space in all models, rendering systems more ‘expressive’ or capable of implementing a broader set of assembly pathways, just like in neural networks. Biologically, diverse subunits are exploited to drive specificity in highly ordered assembly via post-translational modification (34), or chaperone-guided control of subunit binding (35) or disassembly (36). Sequential translation of distinct assembly components (37) or spatial localization further regulate the assembly hierarchy (38). Our AD driven optimization approach enables sampling in such large parameter spaces regardless of internal or external control elements, and thus offers a tool to immediately expand to more complex nonequilibrium protocols.

Far-from-equilibrium protocols are particularly important for evolved systems, as although internal design of binding kinetics provides optimal efficiency ‘for free’ (Fig 5), evolution via gene fusion, for example(39), often produces complexes with repeated subunits and symmetry (40, 41), limiting inherent diversification of binding kinetics. Repeated subunit building blocks also more naturally lends itself for non-monomer growth (23), which we found is always less efficient than monomer growth (Fig 5 and SI), but can still effectively avoid kinetic traps. Such modular growth of stable sub-complexes effectively reduces *N*, and is used by ATP synthetase (42) and the transcription pre-initiation complex, where *N* reduces from 46 to 10 (43).

For designed systems, achieving the bimolecular rate constants of protocol A does present a challenge; predicting rates from molecular interfaces requires theoretical approximations about the free energy barrier between unbound and transition states (44) and expensive (and inexact) computational methods (45). Nonetheless, molecular contacts have been successfully designed for binding kinetics using automatic differentiation (12) and rational design of electrostatic complementarity (46), with 5000-fold differences in rates possible(47). Variation of interface sizes can produce diversification of rates (31), and phosphorylation can be designed to trigger structural changes that promote oligomerization via rate-growth (48). Accessing protein design for diverse asymmetric structures (49) requires understanding the multitude of ways that kinetics can ensure high-yield assemblies, and our study emphasizes the benefits of incorporating kinetics into rational design of protein binding interfaces.

## ACKNOWLEDGEMENTS

M.E.J. gratefully acknowledges funding from a National Institutes of Health MIRA Award R35GM133644. We acknowledge use of the ARCH supercomputer at Johns Hopkins. We thank Dr. Yiben Fu for helping with plotting, and Dr Alex Sodt and Johnson Lab members for feedback.

