## Supplemental Information for "Discovering optimal kinetic pathways for self-assembly using automatic differentiation"

for

### SUPPLEMENTAL METHODS

#### SI. Defining kinetic models of self-assembly

The kinetics of our assembly models are solved using systems of ordinary differential equations (ODEs) in Python, with a few examples also solved using spatial and stochastic particle-based reaction-diffusion models described later. Rather than write out the ODEs for each model, we instead construct them automatically from input files formatted following BioNetGen rules(1) that specify the reaction network. The reaction network specifies all pairwise association ($2^{nd}$ order) and dissociation ($1^{st}$ order) reactions between assembly species, as all binding is reversible. No misinteractions are allowed, thus all intermediates are capable of forming complete complexes. Enzyme reactions are added in the input file as Michaelis-Menten enzymatic reaction, such that the enzyme *E* binds to a complex *AB* to drive its dissociation, $AB + E \rightleftharpoons EAB \rightarrow E + A + B$, with reversible binding rates $k_{\mathrm{f}}$ and $k_{-}$ and catalysis rate $k_{\mathrm{cat}}$. Titration or 'activation' of subunits are modelled as $0^{\mathrm{th}}$ order reactions with rates $\alpha_1$, $\alpha_2$, .. $\alpha_{\mathrm{N}}$ for *N* subunits.

**SII. Numerical Integration Approach and automatic differentiation**

For learning optimal kinetic rates for these biological assembly protocols we use automatic differentiation (AD), which will identify local optima for a target objective function (e.g. yield at a specified time). AD constructs the gradient with respect to our selected parameters (rates) via accumulated computer operations; the advantage over alternate gradient-based methods is it does not require analytical solutions, which do not exist for self-assembly yield, and it does not require the multiple computations needed for the inexact and error-prone numerical gradient(2). The AD method (in reverse mode) offers a generalized form of the backpropagation used in artificial neural networks, and by implementing a differentiable numerical integration of our self-assembly models in PyTorch (3), we can access these efficient routines for high-dimensional parameter spaces.

To exploit the AD functionality in PyTorch(3), we store all variables in our differential equations as torch.Tensor type, and then implement a numerical integration scheme that operates on these datatypes. In this way, we can call on the backward operations of the automatic differentiation package to construct a graph of all variables and operations and every state they existed in over the course of the calculation (in this case the numerical integration of ODEs). With this graph we can compute gradients with respect to our parameters of interest (i.e. rates). We set up our numerical integration scheme using matrix operations to improve efficiency. We use a deterministic event-driven integrator that updates concentrations in fixed intervals rather than a fixed-time-step method like the trapezoid rule. The timestep is chosen at each iteration to achieve a fixed total change in concentration. We found this method to be more effective in balancing accuracy and efficiency of the numerical integration, and provided more stability in gradient calculations given that our changes in concentration were fixed to finite amounts. We validated that this integration scheme produced the correct kinetics by solving the same trimer and tetramer systems using the built-in stiff solvers in Python and MATLAB (e.g. Fig S2). The integration scheme is implemented as follows:

1a) Initialization: We construct the reaction stoichiometry matrix *A* of size *L*x*M* where *L* is the number of species (including all intermediates) and *M* is the total number of reactions including binding and dissociation reactions. If species *i* is a product of reaction *j*, $A_{ij}$=1. If it is a reactant of reaction *j*, $A_{ij}$=-1. Otherwise $A_{ij}$ =0, with factors of 2 for self-interactions.

1b) Initialization: We compute the rate constant matrix in logarithmic form $\ln \vec{k}$. This is a vector of length *M*. The dissociation rates are calculated from the association rates and free energy change of each reaction that breaks *m* bonds by $\ln k_{-,\mathrm{m}} = m\Delta G/(k_{\mathrm{B}}T) + \ln c_0 k_{\mathrm{f}}$ (Eq 2 main text).

2) The instantaneous reaction rates or reaction propensities(4) are evaluated by $\ln \vec{r} = \ln \vec{k} + \ln \vec{X}$. The $\ln \vec{X}$ enforces the law of mass action. $\ln \vec{X}$ is calculated from *A* and $\vec{x}$,

where $A$ is the reaction matrix and $\vec{x}$ is the vector of species concentrations (length $L$). This calculation involves a series of masking, addition, and multiplication steps to maintain differentiability in this vectorized form. For a bimolecular reaction between species 1 and 2, $r_i = k_i x_1 x_2$, whereas for a unimolecular reaction, $r_i = k_i x_1$.

3) The simulation is progressed by taking a small time step that is given by $\Delta t = \frac{C_\delta}{\sum \exp(\ln(r_i))}$. Here $C_\delta$ is a fixed concentration scale defined by the user, which we typically set as $C_\delta = C_{\text{init}}/100$. The time step is therefore not a fixed quantity as the reaction rates change with time. Early on in the simulations, the timestep is short due to rapid changes in species concentrations, whereas at equilibrium, the $\sum r_i \to 0$ and thus $\Delta t \to \infty$, implying correctly that no further change in concentration $C_\delta$ is possible for the final equilibrated system.

4) The change in concentration for all reactions is then $\vec{a} = \exp(\ln(\Delta t) + \ln(\vec{r}))$. Finally the change in concentration for all species is given by $\Delta\vec{x} = A\vec{a}$. All species are updated $\vec{x} = \vec{x} + \Delta\vec{x}$. Current time is updated $t = t + \Delta t$. Steps 2-4 are repeated until the maximum simulation time is reached.

**SIII. Parameter updates for optimization**

Our target objective function we are trying to optimize is the yield at a specified stopping time. The independent parameters we are modifying to maximize yield are rate constants. The yield can be generally defined as $Y = F(\vec{k}, \Delta G, \overrightarrow{x_0}, A, t_{\text{stop}})$, where $t_{\text{stop}}$ is the stopping time and the yield can only be calculated numerically for these coupled nonlinear systems. We choose $t_{\text{stop}}$ such that it is after $\tau_1$ (see Figure 2 Main text), because then it is after trapping sets in, but significantly before $\tau_2$ where the system finally exits from the trapped state under uniform rates. We define a loss function that includes regularization terms ($R$) to ensure the rates remain positive and between our minimum and maximum thresholds, because the optimization can become unstable if parameters are constrained directly. A strong linear penalty is added if rates exceed our min/max thresholds, and the same penalty is used if copy numbers become negative. This penalty will far outweigh the target metric and prevents unphysical solutions. The optimization parameters and loss function for each protocol are described in Table S1.

We normalize the yield $Y$ to its maximum value, which is controlled by the monomer subunit (element of $\vec{x}_N$ where $N$ is number of distinct subunits) with the lowest initial concentration. Each time the numerical integration of the ODEs reaches $t_{\mathrm{stop}}$, the AD method in reverse mode computes the gradients $\frac{\partial L}{\partial k_j}$ for each association rate $k_j$ from the stored graph of partial derivatives. We note that because the free energies are fixed, we do not need gradients with respect to dissociation rates, those are updated from the association rates as defined by Eq. 2. We then update our association rate parameters via $k_j = k_j + \lambda \frac{\partial L}{\partial k_j}$, where $\lambda$ is the learning rate. The convergence to a parameter set that optimized yield at our specified stop time $t_{\mathrm{stop}}$ was dependent on the learning rate, which we chose as fast as possible before numerical instabilities emerged (typically $k_{\mathrm{f}}^{\mathrm{init}}/1000$). This protocol is repeated for $N_{\mathrm{it}}$ iterations until the desired yield is obtained (typically defined by calculating the equilibrium yield in advance).

### SIV. Kinetic Protocols

As introduced in the main text, we use three classes of protocols to restrict which rates constants can be optimized to maximize yield in time. These protocols can be specified by the user in the input file. The optimization parameters varied depending on the kinetic protocol and are described below

*Protocol A:* In this protocol, the parameters are the pairwise binding rates. For the rate growth (A1), we allow only a subset of association rates $\overrightarrow{k_{\mathrm{f}}} = \{k_2, k_3, \dots, k_N\}$ to be optimized. We enforce that the assembly proceeds only by monomer addition/dissociation reactions. We assume the subunits are different species, but the dimerization reactions all occur with the same rate, $k_2$. Similarly, all higher order complexes of size n are constrained to form at a rate $k_n$. For an assembly of size $N$, there are $N$-1 distinct rates that can be optimized. In the diversification model (A2), the parameters are association rates of distinct dimerization reactions ($\overrightarrow{k_{\mathrm{f}}}$). For e.g., the rates for a trimer system are $\overrightarrow{k_{\mathrm{f}}} = \{k_{12}, k_{23}, k_{31}\}$. While in the previous protocol the rates of intermediate formation are independent parameters that are optimized, here these rates are constrained by the corresponding dimer rates depending upon the participating interfaces (See Figure 1 in Main text). For an assembly of size $N$, the optimizable parameters are all the $N$*($N$-1)/2 dimerization reactions. We can restrict assemblies to grow either through monomer addition or allow complementary intermediates to combine. If we allow non-monomer growth, we find orthogonal solutions in the Rate-growth protocol where instead of slowing dimerization, we slow down higher-order reactions to produce a large pool of complementary intermediates and eliminate further elongation. In the Diversification

Model, non-monomer growth is always less efficient because complementary intermediates lack the fast binding interfaces needed for rapid final assembly.

The optimization is started with uniform association rates ($\overrightarrow{k_f} = 1\ \mu M^{-1}s^{-1}$) while keeping $\overrightarrow{\Delta G}$, $\overrightarrow{x_0}$, $t_{stop}$ constant. For optimizing these internal rates, we observed that as long as $t_{stop}$ lies between $\tau_1$ and $\tau_2$ the optimization was successful to achieve high yields and evade traps. Each parameter was updated with the same learning rate, which was constant throughout the optimization. To ensure the rates do not exceed the diffusion limit (~$10^2$ $\mu M^{-1}s^{-1}$) , but with a slightly lower ceiling considering it is protein-protein association(5), we impose a penalty when any of the rates exceed 10 $\mu M^{-1}s^{-1}$ (second term in $R$ in Table S1).

*Protocol B:* In this protocol, all internal association rates are fixed at equal values, thus leading to trapping for stable systems. We then optimize rates at which the *N* monomer species are titrated into the system volume, $\alpha_1$, $\alpha_{2,}$ … $\alpha_N$. We evaluated two modes of adding monomer species into the bulk: single-rate and multi-rate. In single-rate, all subunits have the same titration rate $\alpha_1$, $\alpha_{2,}$ … $\alpha_N \equiv \alpha$, meaning only 1 parameter is optimized for the system. In multi-rate, we optimized the *N*-2 parameters corresponding to the titration rates of *N*-2 species, while the remaining two species are present in bulk.

During optimization, $\overrightarrow{k_f}$, $\overrightarrow{\Delta G}$, $\overrightarrow{x_0}$ remain constant while $t_{stop}$ varied in each iteration as the titration rates were updated. In both modes, a subunit titration is stopped once it reaches the target concentration ($C_{targ}$, which we chose as 100 µM). The simulation continues to $t_{stop}$ beyond this point (max. 1 sec) to allow the completion of assembly. The learning rate for the single-rate mode depended on the initial starting value of the titration rates which was different for different assembly sizes. For multi-rate, we observed that having the same learning rate for all parameters led to a solution where all rates were equal. To aid in obtaining a different solution, we opted for different learning rates for each parameter which were randomized as given in Table S1. We used a learning rate scheduler that decreased each λ by a factor of 0.1 once the yield reached 90% to help achieve convergence. Additionally, to drive optimization towards solution with different rates, we add a penalty for the variance of the titration rates.

*Protocol C:* For the enzymatic control, all internal association rates are fixed at equal values, thus leading to trapping for stable systems. We then optimize the rates of disassembly of a substrate intermediate by an enzyme. There are two different modes of enzyme control that are optimized a) Single substrate and b) Multi substrate. For each enzyme reaction there are 3 parameters that can be optimized, initial enzyme concentration ($E_0$), association rate of substrate binding ($k_1$) and catalysis rate ($k_{cat}$) while keeping the ΔG fixed. So, there are 3 parameters to optimize for case a, and 2*(N-2) + 1 parameters for case b since there are 2 parameters for the rates for each reaction with N-2 reactions, and 1 parameter for the concentration.

The optimization of the single substrate mode is performed in two steps. In step 1, we speed up the kinetics of enzyme association by optimizing the rate of binding $k_1$and initial enzyme concentration $E_0$ that minimizes the substrate (for e.g. an AB dimer) and maximizes the formation of the encounter complex ($EAB$). During this stage $k_{cat}$ is kept constant. In step 2, we optimize the catalysis rate that maximizes the yield of the final complex, while using the optimal values $k_1E_0$ obtained from step 1. For optimization of the multi substrate mode, we use an excess enzyme concentration while maintaining the optimal value of $k_1E_0$ from the single-substrate mode. We optimize the rates of catalysis $\overrightarrow{k_{cat}}$ for disassembly of different substrates while maximizing the yield of the final complex. The learning rate is kept constant for all modes.

**SV. Reaction Diffusion (RD) Simulations**

We further validate the results of our deterministic integration scheme with a stochastic, structure-resolved reaction diffusion simulator NERDDS(6). The assembly kinetics are compared for a hetero-trimer assembly under two different configuration of rates i) Non-optimal where $k_{dim} = k_{tri} = 1\mu M^{-1}s^{-1}$ and ii) Optimal where $k_{dim} = 0.2\ \mu M^{-1}s^{-1};\ k_{tri} = 1\mu M^{-1}s^{-1}$, while $C_{init}$ and $\Delta G$ are fixed at $100\ \mu M$ and $-20k_BT$. The RD simulations are performed with the same macroscopic rate and thermodynamic parameters. For a concentration of 100 µM, we used a cubic box of side 118.41 nm starting with 100 copies of each subunit. Each subunit has two interfaces, one interface per binding partner. Free subunits associate through their binding interfaces and form dimers at a macroscopic association rate of $k_{a,dim} = k_{dim}$, same as in the deterministic simulations. However, during formation of a trimer (loop closure), the deterministic simulations assume a 1-step bimolecular binding ($AB+C \rightarrow ABC_{closed}$) whereas in the RD simulations it occurs in two steps that therefore allows for two pathways of assembly, either ($AB+C_{CtoA} \rightarrow ABC_{open}$ and then $ABC_{open} \rightarrow ABC_{closed}$) or ($AB+C_{CtoB} \rightarrow ABC_{open}$ and $ABC_{open} \rightarrow ABC_{closed}$). To account for this increased number of assembly pathways, the macroscopic association rate of adding a subunit to a dimer was set as $k_{a,tri} = k_{tri}/2$, given that there are two ways for a C to bind to a dimer. The second step occurs at a unimolecular rate defined by $k_{\text{close}} = k_{\text{f}} c_0$, (see ref (6) for derivation) where $c_0$is the standard state 1M and $k_{\text{f}}$ is the association rate for the bimolecular reaction involving those two interfaces. Dissociation of a bond occurs at the pairwise dissociation rate $k_b$, and does not use Eq. 2. However, the long lifetime is preserved because full dissociation requires two unbinding events before rebinding or closure occurs, and rebinding ($k_{\text{close}}$) is fast. A complete description of the parameters is given in Table S2.

**SVI. Equilibrium yield**

For these assembly systems, the equilibrium yield depends on initial concentrations, ΔG values, and the network topology. We solve for the equilibrium via numerical optimization of the coupled algebraic equilibrium between all pairwise interactions, with mass conservation, using nsolve from symPy. This method was relatively slow for larger systems, so we instead used the ODEs at long times to define the final equilibrium.

**SVII. Trimer steady-state in the irreversible limit:**

A useful limiting case is irreversible assembly, where the strength of subunit interactions $\Delta G = -\infty$. Under these conditions, and now using a homo-trimer system, the rate equations can be simplified to:

$$\frac{dx_1}{dt} = -2k_1 x_1^2 - k_2 x_1 x_2 \qquad (S1a)$$

$$\frac{dx_2}{dt} = k_1x_1^2 - k_2x_1x_2 \qquad (S1b)$$

where $x_1, x_2$are the monomer and dimer concentrations respectively and $k_1, k_2$ are the rates of dimerization and trimerization. We can always calculate trimers from mass conservation given monomers and dimers, $x_1 + 2x_2 + 3x_3 = x_{\text{tot}}$. To evaluate the asymptotic behaviour of $x_2$, we define $x_2 = ux_1$, such that $\frac{dx_2}{dx_1} = u + x_1\frac{du}{dx_1}$ and then divide Eq S1b by Eq S1a. Assuming $x_1 \neq 0$, we obtain

$$\frac{dx_2}{dx_1} = \frac{k_1x_1^2 - k_2x_1(ux_1)}{-2k_1x_1^2 - k_2x_1(ux_1)} = \frac{k_1 - k_2u}{-2k_1 - k_2u} \; (S2)$$

and then using our change of variable definition, we have

$$x_1\frac{du}{dx_1} = \frac{-k_2u^2+(k_2-2k_1)u - k_1}{2k_1 + k_2u} \Rightarrow \int \frac{1}{x_1}dx_1 = \int \frac{2k_1 + k_2u}{-k_2u^2+(k_2-2k_1)u - k_1}\, du.$$

After integrating on both sides and simplifying, we can obtain the expression for $x_1$

$$x_1 = e^{\mathcal{C}}e^{\frac{-2k_1 - k_2}{A}tan^{-1}\left(\frac{2k_1+k_2\left(2\frac{x_2}{x_1}-1\right)}{A}\right)}\left(k_1\left(2\frac{x_2}{x_1}+1\right) + k_2\left(\left(\frac{x_2}{x_1}\right)^2 - \frac{x_2}{x_1}\right)\right)^{-\frac{1}{2}} \quad (S3)$$

where $A = \sqrt{-4k_1^2 + 8k_1k_2 - k_2^2}$ , and $\mathcal{C}$ is the integration constant. This transcendental relationship between $x_1$and $x_2$is true for all $t$ and using the initial conditions $x_1(0) = 1$ , $x_2(0) = 0$ we can calculate $\mathcal{C}$ by

$$1 = e^{\mathcal{C}}e^{\frac{-2k_1 - k_2}{A}tan^{-1}\left(\frac{2k_1x_1+k_2(2x_2-x_1)}{Ax_1}\right)}\left(k_1(2x_2x_1 + x_1^2) + k_2({x_2}^2 - x_2x_1)\right)^{\frac{-1}{2}}$$

Thus,

$$e^{\mathcal{C}} = k_1^{\frac{1}{2}}e^{\frac{2k_1 + k_2}{A}tan^{-1}\left(\frac{2k_1 - k_2}{A}\right)} \qquad \text{(S4)}$$

To evaluate the asymptotic value of $x_2$, we evaluate Eq S3 as $x_1 \to 0$ when $t \to \infty$, so

$$tan^{-1}\left(\frac{2k_1x_1 + k_2(2x_2 - x_1)}{Ax_1}\right) \to \frac{\pi}{2}$$

$$\Rightarrow 1 = k_1^{\frac{1}{2}} e^{\frac{2k_1 + k_2}{A} tan^{-1}\left(\frac{2k_1 - k_2}{A}\right)} e^{\frac{(-2k_1 - k_2)\pi}{2A}} k_2^{\frac{-1}{2}} x_2^{-1}$$

$$\Rightarrow x_2 = k_1^{\frac{1}{2}} e^{\frac{2k_1 + k_2}{A} tan^{-1}\left(\frac{2k_1 - k_2}{A}\right)} e^{\frac{(-2k_1 - k_2)\pi}{2A}} k_2^{\frac{-1}{2}} \qquad (S5)$$

Substituting $k_1 = k_2 = 1$, we get the value of $x_2 = e^{\frac{-\pi}{\sqrt{3}}}$. This gives us the trimer yield as 0.674. With equal rates and equal initial concentrations of subunit monomers, the trimer subunits will become trapped in a mixture of 32.6% dimers and 67.4% trimers, independent of the value of the concentration or rate. For a reversible trimer assembly, the yield at trapped state approaches this value as $\Delta G \rightarrow -\infty$. The result is the same for the homo- or hetero-trimer system. The same phenomenon is true for any higher-order cycle-forming assembly (Fig S5). Linear systems will not become trapped for equal stoichiometries (Fig S7).

Eqs S3 and S5 indicate that the fraction of monomers and dimers depends on the relative value of the rate constants. To avoid trapping entirely, we want dimers $x_2 \rightarrow 0$. From Eq. S5, we assume a solution will have a value of $A = \sqrt{-4k_1^2 + 8k_1k_2 - k_2^2}$ that is real. Therefore

$$-4k_1^2 + 8k_1k_2 - k_2^2 \geq 0 \Rightarrow -4r^2 + 8r - 1 \geq 0 \Rightarrow 1 - \frac{\sqrt{3}}{2} < r < 1 + \frac{\sqrt{3}}{2} \text{ (S6)}$$

where $r = \frac{k_1}{k_2}$. From Eq. S6, we can hypothesize that for the system to avoid kinetic trapping and have all monomers converted to trimers, then $\frac{k_1}{k_2} \leq 1 - \frac{\sqrt{3}}{2}$ . Therefore, a ratio of $\frac{k_2}{k_1} \geq 7.46$ will guarantee maximum yield of trimers under irreversible conditions, with fully 100% trimers. This bound is very close to the one we find for 95% trimer yield numerically of $\frac{k_2}{k_1} = 5.8$ ($\Delta G = -100k_\text{B}T$).

**SIX. Scaling of the titration rate with concentration and on-rates**

We briefly describe how an optimal titration rate can be defined based on the kinetics of self-assembly, following the derivation introduced for single-component viral assembly(7). The optimal titration rate should limit the time that a new protein appears in the assembly volume to below the timescale $\tau$ that a protein will bind to a single nucleated structure in its volume $V$. Then each new protein contributes to growth of the single nuclei rather than formation of a distinct one. The titration rate should therefore obey $\alpha^* \leq \frac{1}{\tau V}$. The volume is defined to contain monomers needed for a single assembly, $V = \frac{N}{C_{\text{init}}}$. The timescale for binding of a protein to a single nuclei is defined by mapping to a well-defined problem in the theory of diffusion-influenced reactions for binding to a spherical reactant in a fixed volume (8). For irreversible binding, the key result is that $\tau \propto V/k_{\text{f}}$ over a large range of rates until binding is fully diffusion-limited, and $k_{\text{f}}$ is the rate-limiting subunit-subunit association rate. Thus, the optimal $\alpha^* \propto k_{\text{f}} C_{\text{init}}^2$.

**SX. Differential equations for the trimer model:**

We explicitly define the ODEs for the trimer here:

$$\frac{d[A(t)]}{dt} = -k_f[A(t)][B(t)] - k_f[A(t)][C(t)] - k_f[A(t)][BC(t)] + k_-[AB(t)] + k_-[AC(t)] + k_{-,2}[ABC(t)] \quad \text{(S7a)}$$

$$\frac{d[B(t)]}{dt} = -k_f[A(t)][B(t)] - k_f[C(t)][B(t)] - k_f[B(t)][AC(t)] + k_-[AB(t)] + k_-[BC(t)] + k_{-,2}[ABC(t)] \quad \text{(S7b)}$$

$$\frac{d[C(t)]}{dt} = -k_f[A(t)][C(t)] - k_f[C(t)][B(t)] - k_f[C(t)][AB(t)] + k_-[AC(t)] + k_-[BC(t)] + k_{-,2}[ABC(t)] \quad \text{(S7c)}$$

$$\frac{d[AB(t)]}{dt} = +k_f[A(t)][B(t)] - k_f[C(t)][AB(t)] - k_-[AB(t)] + k_{-,2}[ABC(t)] \quad \text{(S7d)}$$

$$\frac{d[BC(t)]}{dt} = +k_f[C(t)][B(t)] - k_f[A(t)][BC(t)] - k_-[BC(t)] + k_{-,2}[ABC(t)] \quad \text{(S7e)}$$

$$\frac{d[AC(t)]}{dt} = +k_f[A(t)][C(t)] - k_f[B(t)][AC(t)] - k_-[AC(t)] + k_{-,2}[ABC(t)] \quad \text{(S7f)}$$

$$\frac{d[ABC(t)]}{dt} = +k_f[A(t)][BC(t)] + k_f[B(t)][AC(t)] + k_f[C(t)][AB(t)] - 3k_{-,2}[ABC(t)] \quad \text{(S7g)}$$

These equations ensure the three mass conservation relations, $C_{\text{init}} = [A(t)] + [AB(t)] + [AC(t)] + [ABC(t)]$, $C_{\text{init}} = [B(t)] + [AB(t)] + [BC(t)] + [ABC(t)]$, and $C_{\text{init}} = [C(t)] + [AC(t)] + [BC(t)] + [ABC(t)]$.

**SXI. Renormalizing variables in the uniform parameter systems to illustrate TF independence on changes to equal rates**

Our systems of differential equations for our self-assembly models are all autonomous, or without any explicit time variables. Thus, if we define a new time $t^* = k_{\text{f}}C_{\text{init}}t$, then $\frac{d[A]}{dt} = k_{\text{f}}C_{\text{init}}\frac{d[A]}{dt^*}$ and all systems with uniform rates become dependent only on the ratios $\frac{k_{-,m}}{k_f}$. If we define all concentrations $[X(t)]^* = [X(t)]/C_{\text{init}}$, then the coefficient for all association reactions becomes 1 and for dissociation reactions the coefficients are $K_D/C_{\text{init}}\exp((m-1)\Delta G/k_{\text{B}}T)$, where m are the edges that break for this intermediate to dissociate a monomer (plus any stoichiometric factors).

e.g. for the trimer system defined in Eq. S7, Eq S7a becomes:

$$\frac{d[A(t)]^*}{dt^*} = -[A(t)]^*[B(t)]^* - [A(t)]^*[C(t)]^* - [A(t)]^*[BC(t)]^* + \frac{K_{\text{D}}}{C_{\text{init}}}[AB(t)]^* + \frac{K_{\text{D}}}{C_{\text{init}}}[AC(t)]^* + \frac{K_{\text{D}}}{C_{\text{init}}}\exp(\Delta G/k_{\text{B}}T)[ABC(t)]^*, \quad \text{(S8)}$$

with all initial conditions now dimensionless values of 1 for monomers or otherwise zero, and the similar structure for all other variables. If we alternately use the same concentration rescaling by choose $t' = k\_t$, an equivalent form for Eq S7a becomes:

$$\frac{d[A(t)]^*}{dt'} = -\frac{C_{\text{init}}}{K_{\text{D}}}[A(t)]^*[B(t)]^* - \frac{C_{\text{init}}}{K_{\text{D}}}[A(t)]^*[C(t)]^* - \frac{C_{\text{init}}}{K_{\text{D}}}[A(t)]^*[BC(t)]^* + [AB(t)]^* + [AC(t)]^* + \exp(\Delta G/k_{\text{B}}T)[ABC(t)]^*, \qquad \text{(S9)}$$

While the absolute time and therefore efficiency depends on the rates via $t^* = k_{\text{f}}C_{\text{init}}t$, any ratio of timescales such as the TF will depend only on the parameters $\Delta G$ and $C_{init}$ based on Eq. S8 or S9. Therefore, if we keep $\Delta G$ and $C_{\text{init}}$ fixed but speed up the on and off-rates, the timescales will be shorter but the TF will not change (Fig S6).

### SUPPLEMENTAL TABLES

**Table S1:** Optimization Parameters and objective functions for each protocol.

| Protocol | Yield | Regularization | Loss Function | Learning rate |
|---|---|---|---|---|
| A | $Y = \frac{[Final\ Complex]}{\min_{x_0 \in \vec{x}_N} x_0}$ <br> $t_{stop} = 1\ s$ | $R = ReLU(10 * \lambda - \overrightarrow{k_f}) + ReLU(\overrightarrow{k_f} - 10)$ | $L = -Y + R$ <br> $Minimize\ L\ w.r.t\ \overrightarrow{k_f}$ | $\lambda = 0.01$ |
| B <br> (Single rate) | $Y = \frac{[Final\ Complex]}{C_{targ}}$ <br> $t_{stop} = \frac{C_{targ}}{\alpha} + 1\ s$ | $R = ReLU(10 * \lambda - \alpha)$ | $L = -Y + R$ <br> $Minimize\ L\ w.r.t\ \alpha$ | $\lambda = 0.1 * \alpha_0$ <br> $\alpha_0 - Initial\ value$ |

| | | | | |
|---|---|---|---|---|
| B<br>(Multi rate) | $Y = \frac{[Final\ Complex]}{C_{targ}}$<br>$t_{stop} = \max_{\alpha \epsilon \alpha_{N-2}} \{\frac{C_{targ}}{\vec{\alpha}}\} + 1\ s$ | $R = ReLU(10 * \lambda - \vec{\alpha}) +$<br>$ReLU(-1 * Var(\vec{\alpha}))$ | $L = -Y + R$<br>$Minimize\ L\ w.r.t\ \vec{\alpha}$ | $\lambda_i = rand(0.001, 0.1) * \alpha_i$ |
| C<br>(Single Substrate) | Step 1) $S = \frac{[Substrate]}{\min_{x_0 \in \vec{x}_N} x_0}$<br>Step 2) $Y = \frac{[Final\ Complex]}{\min_{x_0 \in \vec{x}_N} x_0}$ | Step 1) $R = ReLU(10 * \lambda - k_1) + ReLU(E_0 - 1)$<br>Step 2) $R = ReLU(10 * \lambda - k_{cat})$ | Step 1) $L = -S + R$<br>$Minimize\ L\ w.r.t\ k_1\ and\ E_0$<br>Step 2) $L = -Y + R$<br>$Minimize\ L\ w.r.t.k_{cat}$ | Step 1) $\lambda = 0.01\ (k_1); 1\ (E_0)$<br>Step 2) $\lambda = 0.001$ |
| C<br>(Multi Substrate) | $Y = \frac{[Final\ Complex]}{\min_{x_0 \in \vec{x}_N} x_0}$ | $R = ReLU\big(10 * \lambda - \overrightarrow{k_{cat}}\big)$ | $L = -Y + R$<br>$Minimize\ L\ w.r.t.\overrightarrow{k_{cat}}$ | $\lambda = 0.001$ |

**Table S2:** Simulation parameters for the RD simulations

| Parameter | Description | Value |
|---|---|---|
| **Simulation details:** | | |
| $\Delta t$<br>Box size | Simulation time step<br>Length of box edge | $0.3\ \mu s$<br>$118.41\ nm$ |
| **Molecule Parameters:** | | |
| $D_t$<br>$D_r$<br>checkOverlap<br>overlapSepLimit | Translational Diffusion constant<br>Rotational Diffusion constant<br>Associations resulting in steric overlap are prevented | $22\ \mu m^2/s$<br>$0.17\ rad^2/\mu s$<br>True |
| **Reaction Parameters:** | | |

| | | |
|---|---|---|
| $k_{a,dim}$ | Macroscopic binding rate for dimerization | $1\ \mu M^{-1}s^{-1}\ (trapped)$; $0.2\ \mu M^{-1}s^{-1}(optimal)$ |
| $k_{a,tri}$ | Macroscopic binding rate for trimerization | $0.5\ \mu M^{-1}s^{-1}$ |
| $k_b$ | Microscopic dissociation rate | $0.00206\ s^{-1}$ |
| $\sigma$ | Distance between two reacting interfaces | $1.0\ nm$ |
| loopCoopfactor | Scaling factor multiplied to $k_{a,tri}$during loop closure | $1.0$ |

**Table S3:** Comparison of $\Delta G_{opt}$ evaluated from exponential fit of main text Eq 3b and found numerically by solving the differential equations (ODEs) and finding where the timescale $\tau_{95}$ is fastest. $C_{init}$ = 100 μM.

| **Assembly Size** | **Fitted $\Delta G_{opt}$** | **ODE based $\Delta G_{opt}$** |
|---|---|---|
| 3 | -10.67 $k_BT$ | -11.5 $k_BT$ |
| 4 | -7.06 $k_BT$ | -8.3 $k_BT$ |
| 5 | -4.65 $k_BT$ | -7 $k_BT$ |
| 6 | -4.35 $k_BT$ | -6 $k_BT$ |
| 7 | -4.01 $k_BT$ | -5.55 $k_BT$ |

**SUPPLEMENTAL FIGURES**

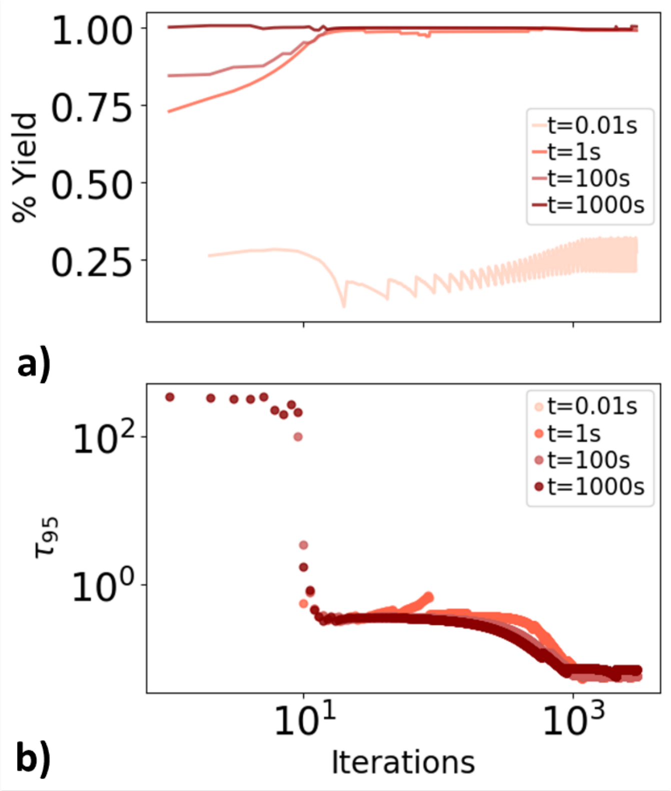


**Figure S1: Optimization of trimer assembly for different stop times ($t_{stop}$).** a) If $t_{stop}$ is too early (0.01s beige curve), the system is physically incapable of reaching its equilibrium yield within that time, given that we have a maximal threshold for $k_f$. We must choose $t_{stop}>\tau_1$, where $\tau_1$ is where defined as an inflection point in the yield kinetics vs ln(t). For $t_{stop}>\tau_1$, we can achieve maximal yield via rate optimization, as illustrated here for 3 values (1, 100, and 1000s). For t=1000s, the maximal yield is practically already reached by this time, but the optimization still accelerates the rates, because it gives earlier timepoints better yield, thus marginally improving the yield at longer times. b) The timescale (unnormalized) $\tau_{95}$ as we iterate over new solutions improves systematically. If the yield does not reach 95%, data points are not shown. The learning rate used is 0.01 for all simulations.

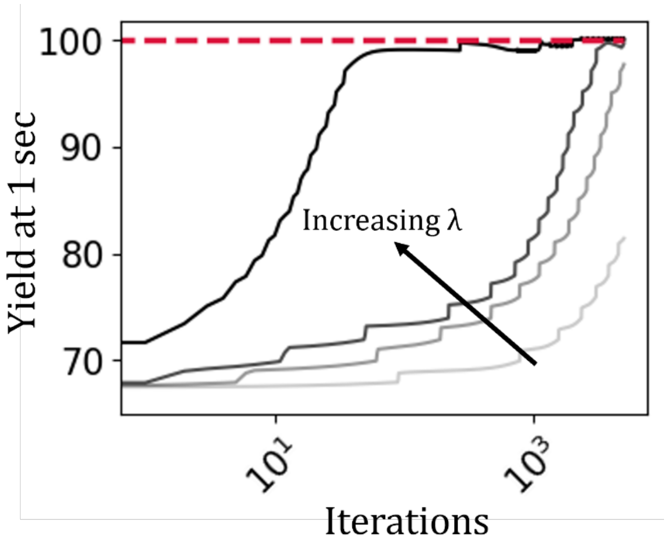


**Figure S2: Optimization of trimer assembly with different learning rates (λ).** The optimization is performed with different learning rates (increasing from light grey to dark gray: 1e$^{-4}$, 5e$^{-4}$, 1e$^{-3}$, 5e$^{-3}$) for 5000 iterations. All optimizations are started with the initial rate parameters $\overrightarrow{k_{f,0}} = 1\ \mu M^{-1}s^{-1}$. The choice of learning rates affects convergence of model parameters to more optimal values. For optimization of most of the protocols setting $\lambda = (1e^{-2} - 1e^{-3}) * \overrightarrow{k_{f,0}}$ helped achieve convergence quickly.

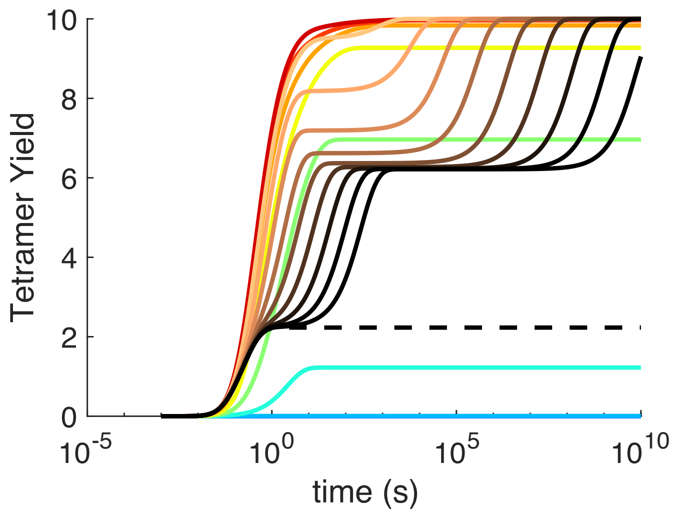


**Figure S3. Tetramer yield vs time with increasing free energy Δ*G*.** All curves have a fixed initial concentration for each subunit $C_{init}$=10$\mu$M, and a fixed $k_f$=1$\mu$M$^{-1}$s$^{-1}$ for all binding events. Monomer growth only. For the blue to dark red curves, the free energy is increasing from -1$k_BT$ to -11 $k_BT$ , and the yield is continually improving. With yield <99.9%, we do not have trapping, and the TF=0. For the light to dark brown curves, the TF>0 and increasing as we continue to increase ΔG from -12 $k_BT$ to -20 $k_BT$ . The black dashed curve is the kinetics for fully irreversible tetramer assembly Δ*G*= - **∞.** For the tetramer, we see the clear emergence of multiple plateau regions, with the first plateau being the delay due to dimer dissociation, and the second plateau being the longer delay due to trimer dissociation. The TF is always defined based on the first entry time $\tau_1$ and the *final* exit time $\tau_2$ from trapping, ignoring all intermediate plateaus. Here $\tau_1$~ 0.1s given the fixed $C_{\text{init}}k_{\text{f}}$, and $\tau_2$ keeps lengthening as Δ*G* increases, reaching ~5x10$^9$s for -20 $k_{\text{B}}T$. The most efficient solution is where the yield first reaches ~99.9% and $\tau_{95}$~7s. There is then a regime -12<ΔG<-11 $k_{\text{B}}T$ where the efficiency drops but we still measure TF=0 based on our definition requiring at least two distinct maxima in the derivative d*Yield*/dln(*t*).

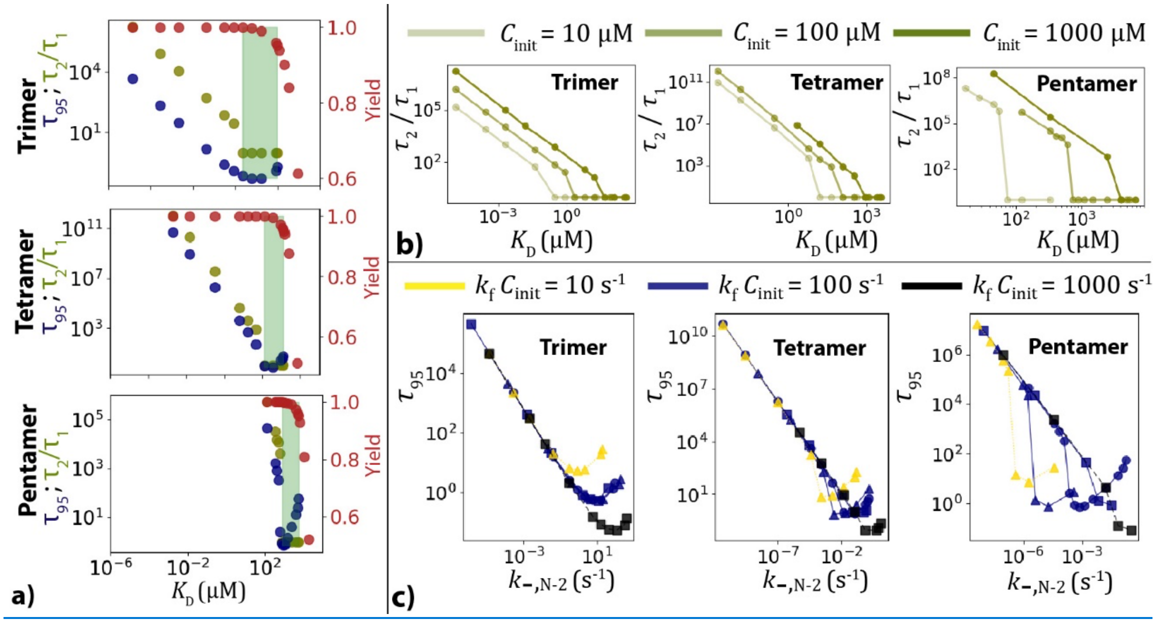


**Figure S4. Timescales of trapping versus interaction strength (Δ*G*) and initial concentration $C_{\text{init}}$.** a) Onset of trapping (TF>1) coincides with the yield reaching >99%. We define a *ΔG* ($K_D$/1M=exp(*ΔG*/$k_BT$)) window shown as the shaded green region where yield is >95% and the TF =1. For trimer, tetramer and pentamer at an initial concentration of 100 μM the time to reach 95% yield $\tau_{95}$ scales the same way as the TF in the trapped regime and reaches its fastest value in the shaded green region, defining $\Delta G_{\text{opt}}$. b) TF evaluated at 3 different initial concentrations of 10 μM, 100 μM, 1000 μM for trimer, tetramer and pentamer. Concentration does not change the scaling with $K_D$ just the magnitude of TF. c) $\tau_{95}$ for different rates and concentrations collapses to an asymptotic universal scaling vs the slowest dissociation rate, corresponding to the *N*-1 size intermediate which has *m*=*N*-2 bonds, or $k_{-,N-2}$ (Eq 2).

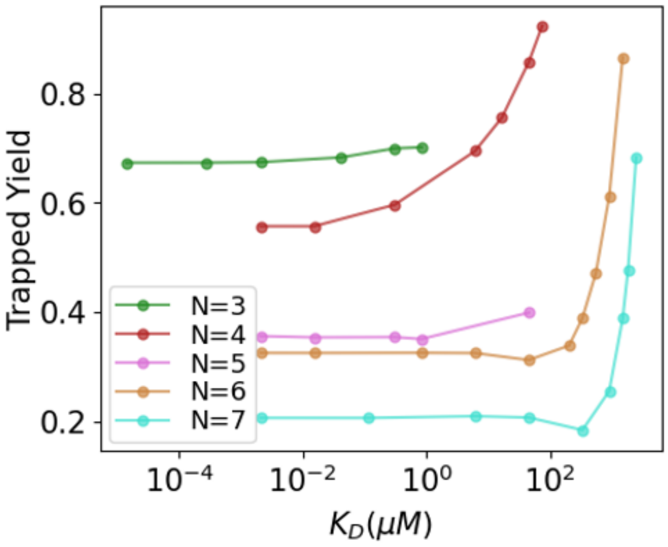


**Figure S5: Yield of the assemblies of different sizes at the trapped state.** As the strength of subunit interactions increases, the fraction of monomers present in *N*-1 intermediate starts to increase which drops the yield of the final complex at the trapped state. As the trapped yield plateaus, the monomers are distributed amongst more sizes of intermediates, which contributes to the increasing TF. These models include monomer and non-monomer growth, hence the improved yield in the trapped state for the tetramer relative to Fig S3. With only monomer growth allowed, the trapped yield will drop for all values of *N.*

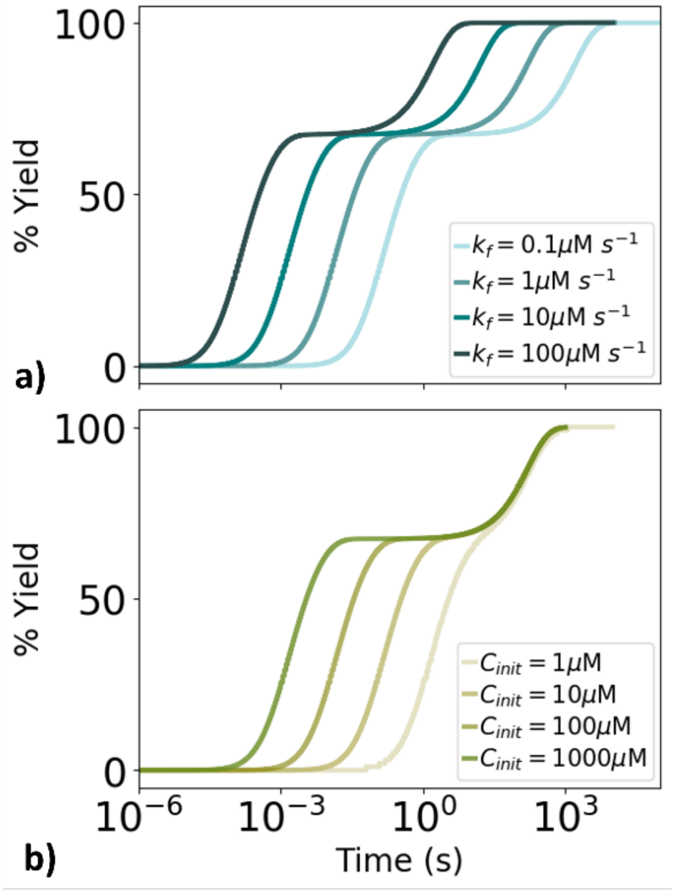


**Figure S6: Concentration profile of trimer kinetics under different binding rates $k_f$ (constant $\Delta G$, $C_{init}$ ) and concentrations $C_{init}$ (constant $\Delta G$, $k_f$).** a) Accelerating the forward/reverse rates while keeping the $\Delta G$ fixed does not eliminate trapping, only shifts the timescales of entry and exit. $C_{init}$=100$\mu$M, $\Delta G$=-20$k_B T$ and %Yield=[Trimers]/$C_{init}$*100. b) Increasing the initial concentrations speeds up the entry into the trapped state while the exit times remain unchanged with $k_f$ =1 $\mu M^{-1} s^{-1}$, and $\Delta G$=-20$k_B T$.

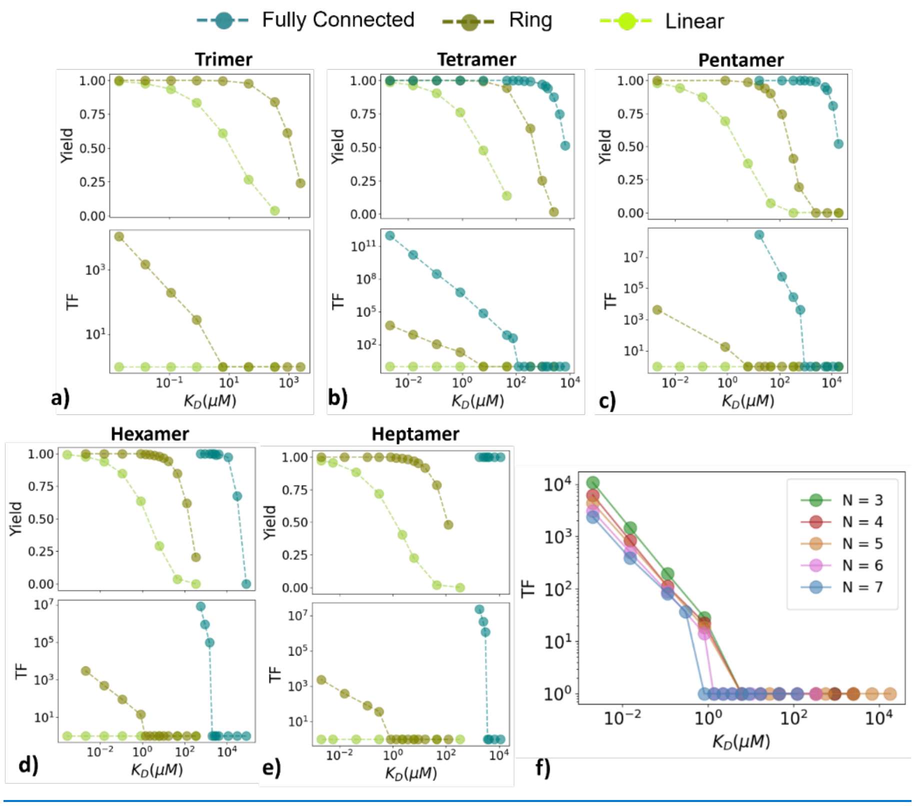


**Figure S7: Yield and trapping factor for distinct topologies.** (a-e) For the linear (light green) and ring (olive green) topologies, as the number of subunits grows from *N*=3 (a) to *N*=7 (e), the interactions must become more stable (lower $K_D$) to achieve a high yield (upper plots). In contrast, the fully connected topology (blue) reaches high yield with weaker contacts as more subunits are added, due to the increase in total binding contacts per subunit. The fully connected topology also has a much steeper dependence of yield on $K_D$ as the number of subunits increases. The fully connected topology suffers much more in terms of timescales once kinetic trapping sets in (lower plots). Linear topologies do not get trapped with these equal stoichiometries. The concentrations used are $C_{init}$=100$\mu$M, and the on rates are $k_f$ = 1$\mu M^{-1}s^{-1}$. (f)The extent of trapping does not get worse with assembly size for the ring topology, unlike in the fully connected case.

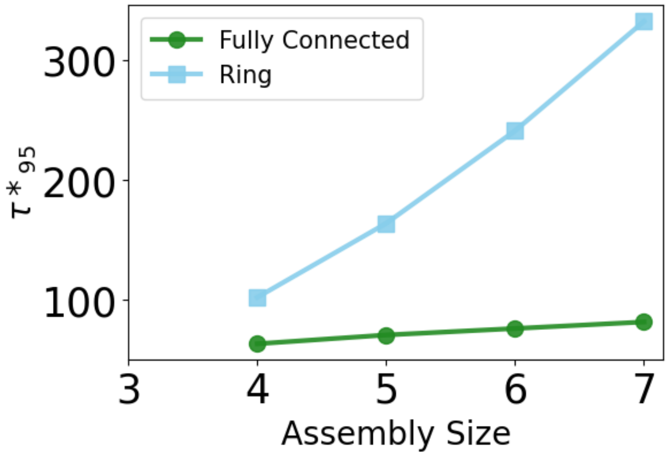


**Figure S8: Efficiency of two topologies at their corresponding $\Delta G_{opt}$ values.** The ring assembly has significantly lower efficiency than the fully connected topology for the optimal reversible assembly at their $\Delta G_{opt}$ values. This is because of the faster dissociation of all intermediates in the ring topology that proceed at the same $k_-$, whereas for the fully connected topology $k_{-,m}$ slows significantly for larger intermediates. The simulations were performed with $k_f$ = 1μ$M^{-1}$ $s^{-1}$ and $C_{init}$ = 100μM and ΔG = $\Delta G_{opt}$ which for the ring topology is -12$k_BT$ for N=4, -12.5 $k_BT$ for N=5, -12.8 $k_BT$ for N=6 and –13.3 $k_BT$ for N=7.

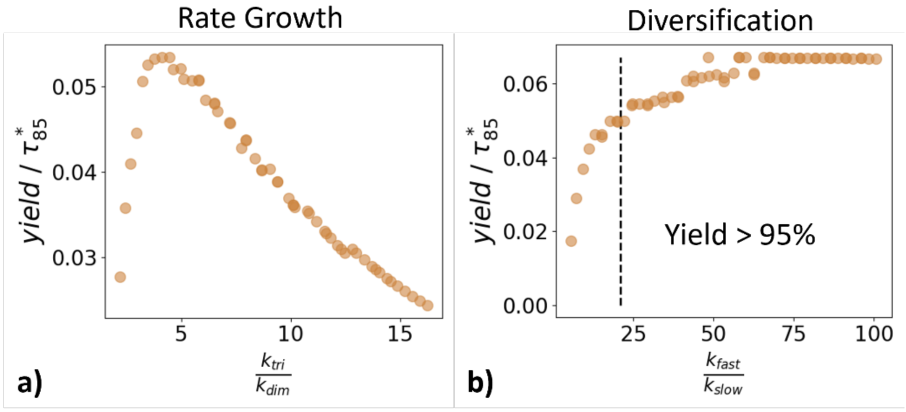


**Figure S9: Separation of rates during optimization for protocol A for a heterotrimer**. The optimization was performed with a learning rate of 0.01 and $t_{stop}$ =1 sec for 5000 iterations. The modified rates are evaluated as ratios - a) $k_{tri}/k_{dim}$ for the rate-growth and b) $k_{fast}/k_{slow}$ for dimer diversification and plotted with respect to the efficiency of the assembly which is quantified as the yield per time. Here the yield is taken as 85% and the time is the normalized time it takes to reach this yield. a) In the rate growth model, there exists an optimal ratio between rate of trimerization and dimerization that leads to maximum efficiency. We identified this ratio for different assembly sizes with different interaction strengths (Main text Figure 3). b) For the diversification model, the efficiency increases as the separation increases until it reaches a plateau. To evaluate the optimal solutions for different assembly size and designability, we quantify the minimum separation required to reach 95% yield (Main text Figure 3).

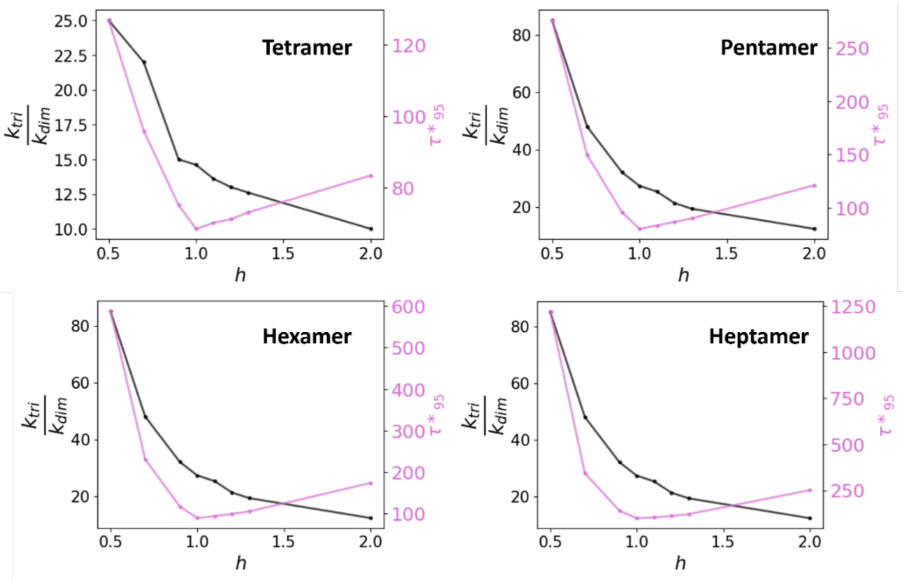


**Figure S10: Ratio of trimerization rate to dimerization rate dependence on speed of subsequent elongation steps for rate-growth model**. The optimal separation between trimerization and dimerization is measured (black line) while fixing the separation between elongation steps ($k_{tetra}, k_{penta}, k_{hexa}, k_{hepta}$) by a factor *h*. For e.g., $k_{tetra} = hk_{tri}, k_{penta} = hk_{tetra}, k_{hexa} = hk_{penta}, k_{hepta} = hk_{hexa}$. When each step of the elongation increases, the optimal separation required between trimerization and dimerization decreases. Therefore, by introducing additional hierarchy in the elongation steps, we can evade kinetic traps by having a slower rate of trimerization thus aiding designability. However, while having faster elongation steps decreases the time taken to complete assembly, the normalized speed slows ($\tau^*_{95}$) (pink curve), as we always normalize by the fastest rate, which controls fewer reactions. The assembly is most efficient when the rates of subsequent elongation steps are as fast as the trimerization step (*h* = 1).

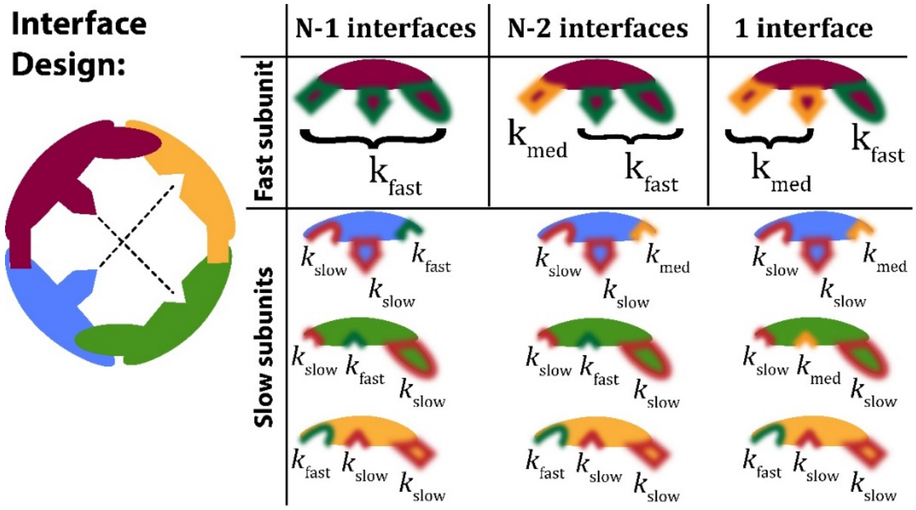


**Figure S11: Designability of interfaces in the Diversification model.** For the fully connected tetramer, we illustrate how our diversification model creates a hierarchy of rate constants for distinct interfaces. The 'best' solution is to accelerate all *N*-1 interfaces on a single subunit. Each of the remaining subunits then has a single 'fast' interface that binds to the red subunit and two slow interfaces. To ease designability, we can also optimize solutions where fewer interfaces have to reach fast binding kinetics. For *N*-2, we then allow a further hierarchy of rates, with a medium speed interface on the otherwise fast-binding subunit. Remaining subunits each still have two slow interfaces. For a single fast interface in the last column, we discover a similar hierarchy of three rates at fast, medium and slow, with each subunit each having at least one fast or medium rate constant. As we discuss in the main text, in this 1-interface model the efficiency is lower, and the ratio between the fast and slow rates must be larger to eliminate trapping. Hence fewer interfaces must be fast, but there must be a larger speed difference to avoid kinetic trapping.

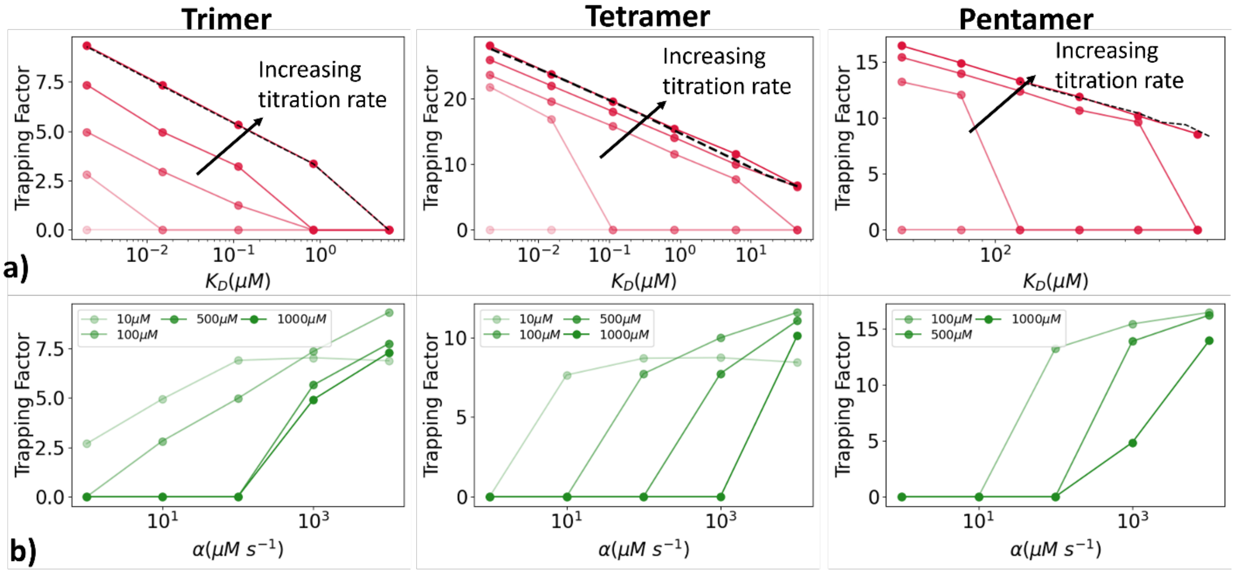


**Figure S12: Extent of trapping under titration for trimer and pentamer.** a) The TF was evaluated with different titration rates ranging from 1 µM s$^{-1}$ (light red) to 10000 µM s$^{-1}$ (dark red) and interaction strengths for trimer to pentamer under a constant target concentration of 100 µM. As seen in bulk, TF not only increases with stability, but also with faster titration rates until it converges to the values observed in bulk conditions (black dashed). b) Increasing the target concentration can allow for a faster titration rate without trapping. This is because assembly proceeds more quickly for higher concentrations (Fig S6), and thus monomers can be similarly titrated in more quickly to complete nascent assemblies.

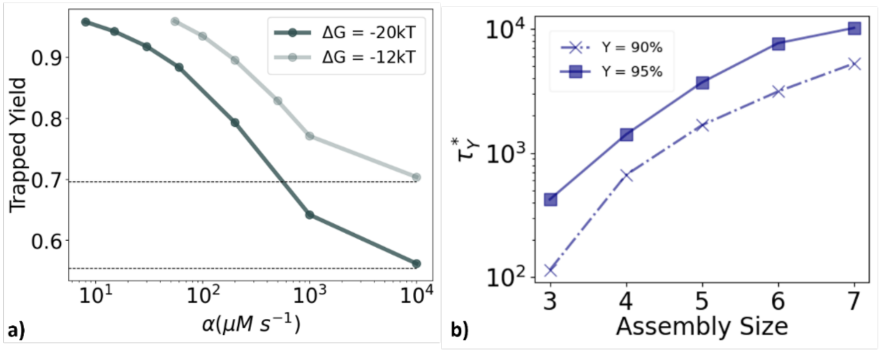


**Figure S13: Efficiency of titration depends upon choice of yield threshold.** a) With faster dissociation of dimers (light grey $\Delta G$=-12$k_BT$) trapped systems reach higher yield. The improvement in yield at slower titration (low $\alpha$) becomes less significant while the drop in efficiency is more significant. b) The efficiency improves when we accept titration protocols that achieve a lower yield threshold of only 90% (dashed x) rather than requiring 95% yield (solid squares). This distinguishes titration protocols from the internally optimized rates of protocol A, where efficiency is very similar for all high-yield (90 vs 95% yield) solutions.

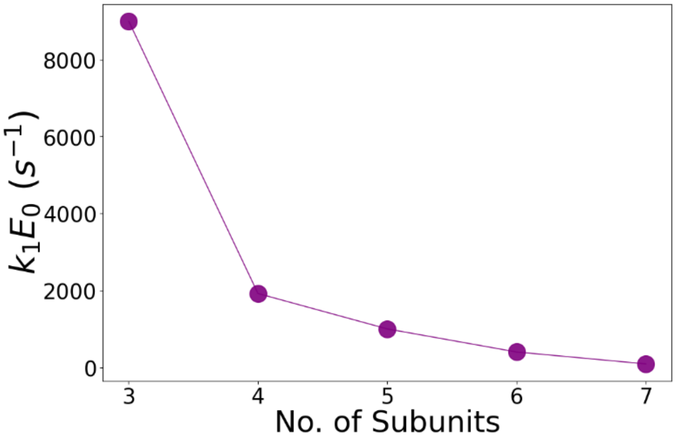


**Figure S14: Optimal values of rate of enzyme binding with its substrate (single) for different assembly sizes.** The optimal rate of enzyme binding as calculated through step 1 of optimization for the single substrate (See Table S1) decreases with assembly size since higher intermediates have lesser competition from their corresponding binding partners. For step 2 of the optimization and for multi-substrate protocol, we fix the enzyme binding rates to these optimal values and optimize the rate of catalysis.

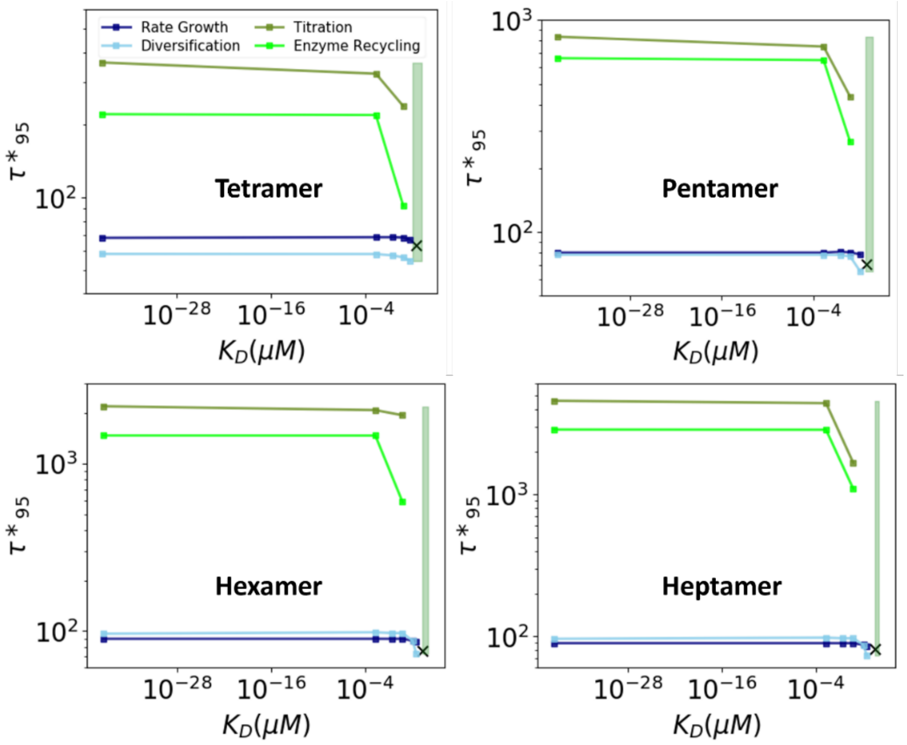

**Figure S15: Efficiency of all protocols optimized at different interaction strengths for each assembly size.** The efficiency of optimized solutions plateaus as the assemblies approach the irreversible limit. Optimization of internal rates (protocol A) are the most efficient methods (blue and light blue) when compared to external protocols (B-olive, C-neon green). The efficiency is improved a small degree for the diversification model when the system has faster dissociation. For the external protocols (green curves), there is a more obvious advantage to optimizing kinetics when the system has faster dissociation. The green shaded region represents the optimal $\Delta G$ window as described in Figure 2 (Main text) and 'X' represents the efficiency obtained when the interaction strengths are optimal ($\Delta G_{opt}$). $C_{\text{init}}$=100$\mu$M for all subunits.
